# The Raf/LIN-45 C-terminal distal tail segment negatively regulates signaling in *Caenorhabditis elegans*

**DOI:** 10.1101/2024.07.16.603803

**Authors:** Robert A. Townley, Kennedy S. Stacy, Fatemeh Cheraghi, Claire C. de la Cova

## Abstract

Raf protein kinases act as Ras-GTP sensing components of the ERK signal transduction pathway in animal cells, influencing cell proliferation, differentiation, and survival. In humans, somatic and germline mutations in the genes *BRAF* and *RAF1* are associated with malignancies and developmental disorders. Recent studies shed light on the structure of activated Raf, a heterotetramer consisting of Raf and 14-3-3 dimers, and raised the possibility that a Raf C-terminal distal tail segment (DTS) regulates activation. We investigated the role of the DTS using the *Caenorhabditis elegans,* which has a single Raf ortholog termed *lin-45*. We discovered that truncations removing the DTS strongly enhanced *lin-45(S312A)*, a weak gain-of-function allele equivalent to *RAF1* mutations found in patients with Noonan Syndrome. We generated mutations to test three elements of the LIN-45 DTS, which we termed the active site binding sequence (ASBS), the KTP motif, and the aromatic cluster. In the context of *lin-45(S312A),* mutation of either the ASBS, KTP motif, or aromatic cluster enhanced activity. We used AlphaFold to predict DTS protein interactions for LIN-45, fly Raf, and human BRAF, within the activated heterotetramer complex. We propose distinct functions for the LIN-45 DTS elements: i) the ASBS binds the kinase active site as an inhibitor, ii) phosphorylation of the KTP motif modulates DTS-kinase domain interaction, and iii) the aromatic cluster anchors the DTS in an inhibitory conformation. This work establishes that the Raf/LIN-45 DTS negatively regulates signaling in *C. elegans* and provides a model for its function in other Raf proteins.

## Introduction

Raf protein kinases are multi-domain signal transduction proteins that act as effectors of GTP-bound Ras, and in turn activate the kinases MEK and ERK to influence cellular differentiation, division, and survival (LAVOIE *et al*. 2020). Humans have three genes encoding Raf, *ARAF, BRAF*, and *RAF1*. Somatic mutations in *BRAF* can occur in malignancies; these most commonly affect the activation loop of the kinase domain and are strongly activating (DAVIES *et al*. 2002). By contrast, inherited and de novo mutations of the *BRAF* or *RAF1* genes are associated with the RASopathy disorders Cardiofaciocutaneous syndrome (CFC) and Noonan Syndrome (NS) respectively, and typically alter the non-catalytic regulatory domains (HEBRON *et al*. 2022).

Vertebrate and invertebrate Raf proteins share three conserved regions (Fig. 1a). Conserved region 1 (CR1) encodes two structural domains: the Ras binding domain (RBD) and cysteine rich domain (CRD), which are both capable of binding Ras-GTP when Raf is active or 14-3-3 when Raf is inactive (GHOSH *et al*. 1994; PARK *et al*. 2019; TRAN *et al*. 2021; MARTINEZ FIESCO *et al*. 2022). Conserved region 2 (CR2) is a short linear motif that constitutes a phospho-dependent binding site for 14-3-3. Conserved region 3 (CR3) is a kinase domain. Additionally, a fourth C-terminal tail region contains a second, highly conserved phospho-dependent 14-3-3 binding site followed by a sequence recently named the distal tail segment (DTS) (KONDO *et al*. 2019; LIAU *et al*. 2020a)

**Figure 1.**
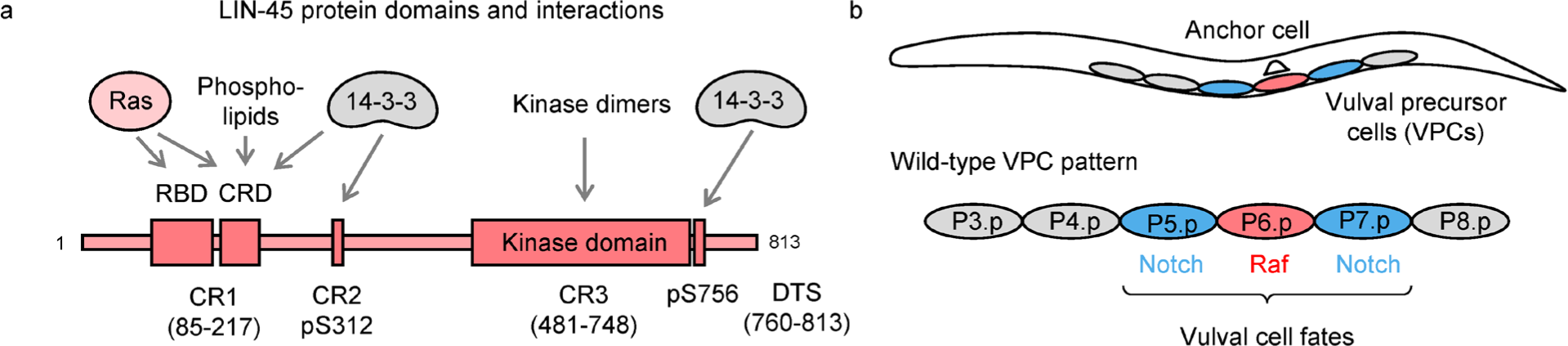
LIN-45 protein domains and activity in *C. elegans* development. **a)** Map of conserved regions in LIN-45. Conserved region 1 (CR1) contains the Ras-binding domain (RBD) and cysteine rich domain (CRD). 14-3-3 binds at phosphorylated S312 of conserved region 2 (CR2) and at phosphorylated S756 of the C-terminal region. Conserved region 3 (CR3) is the kinase domain. The distal tail segment (DTS) consists of sequence between the 14-3-3 binding site and the terminus, residues 760-813. **b)** At the L3 larval stage, EGFR signaling in P6.p leads to Raf activation and 1° fate (red), while Notch activation in P5.p and P7.p results in 2° fate (blue).

The last decade has produced a wealth of structural studies that shed light on the conformation of human BRAF before and during activation (KONDO *et al*. 2019; PARK *et al*. 2019; KHAN *et al*. 2020; KONDO *et al*. 2021; MARTINEZ FIESCO *et al*. 2022). In the absence of Ras-GTP, BRAF is in an off state, phosphorylated at the CR2 and C-terminal 14-3-3 binding sites, permitting each site to be bound by one protomer of a 14-3-3 dimer. In this inactive monomer conformation, the kinase domain is bound to MEK in an inhibitory manner that prevents dimerization and activation. When Ras-GTP is present, it binds directly to the BRAF RBD, causing relocalization to the plasma membrane where the CR2 site is dephosphorylated by a protein complex containing PP1C and SHOC2 (HAUSEMAN *et al*. 2022; LIAU *et al*. 2022). Dephosphorylation is coupled to interaction with KSR, which displaces MEK and promotes formation of Raf homodimers, Raf auto-phosphorylation and activation. In the on state, a BRAF dimer and 14-3-3 dimer form a heterotetramer, where each protomer of BRAF contacts a 14-3-3 protomer through the phosphorylated C-terminal 14-3-3 binding site (PARK *et al*. 2019).

Activation of BRAF ultimately leads to negative feedback when it is directly phosphorylated by ERK at three functionally distinct negative regulatory sites (RITT *et al*. 2010). The first of these is in the RBD, and phosphorylation reduces BRAF-Ras binding. A second phosphorylation is within a Cdc4-phosphodegron (CPD) site and promotes protein degradation (DE LA COVA AND GREENWALD 2012; SAEI *et al*. 2018). A third is within the C-terminal DTS, at residues S750 and T753, where phosphorylation reduces BRAF dimerization (RITT *et al*. 2010). Because these modifications are catalyzed by the downstream kinase ERK, they occur a short time after BRAF activation, causing BRAF activation to transiently peak before declining and returning to baseline. Such feedback is a possible mechanism underlying the dynamic nature of ERK signaling, which can be transient, sustained, or pulsatile in many cell types, including invertebrates (DE LA COVA *et al*. 2017; NAKAMURA *et al*. 2021).

Because most cryo-EM structures of active BRAF dimers do not resolve the C-terminal BRAF DTS region phosphorylated at S750 and T753, little is known about the mechanism of feedback through this site (PARK *et al*. 2019; MARTINEZ FIESCO *et al*. 2022). In the cryo-EM structure that successfully resolves a portion of the DTS, the kinase domains adopt an asymmetric conformation, rotated so that one kinase protomer is closer to the 14-3-3 dimer (KONDO *et al*. 2019). Residues F743-S750 of BRAF were sufficiently ordered to resolve side chains, allowing identification of this segment as occupying the ATP-binding pocket of the kinase active site (KONDO *et al*. 2019). We refer to this sequence throughout this manuscript as the active site binding sequence (ASBS). The BRAF DTS was initially proposed to positively regulate signaling; however, studies directly testing the effect of DTS truncation failed to show signaling differences (KONDO *et al*. 2019; LIAU *et al*. 2020a). A possible obstacle to testing the role of the BRAF DTS is that mammalian cells may express *ARAF*, *BRAF*, and *RAF1* genes, and manipulations that inhibit one Raf can result in paradoxical activation of the pathway (HEIDORN *et al*. 2010; POULIKAKOS *et al*. 2010)

The nematode *Caenorhabditis elegans* has a single Raf ortholog, termed LIN-45, that is required for larval viability, vulval fate specification, germline morphology, meiosis, and oocyte maturation (HAN *et al*. 1993; HSU *et al*. 2002). Phenotypes caused by *lin-45* loss-of-function and gain-of-function during the process of vulval fate specification in larval development are easily observed later in adulthood. During the L2 and L3 stages of larval development, Vulval Precursor Cells (VPCs) are patterned when the Anchor Cell (AC) of the gonad produces an EGF-like ligand (SCHINDLER AND SHERWOOD 2013; SHIN AND REINER 2018). In normal development, signaling by the EGF receptor, Raf, MEK, and ERK is activated in a single VPC termed P6.p. As a consequence, this cell adopts a fate termed “1°,” expresses a Delta/Serrate/LAG-2 (DSL) ligand, and eventually produces vulval structures in the adult. The DSL ligand expressed by P6.p activates Notch signaling in adjacent VPCs, termed P5.p and P7.p, which adopt a vulval fate termed “2°,” while other VPCs that receive no EGF or Notch stimulus adopt a non-vulval fate. Mutations that ablate *lin-45* activity cause failure in vulval fate specification, and a phenotype termed Vulvaless (Vul). Conversely, mutations that increase *lin-45* activity cause more than one VPC to adopt 1° fate, and a phenotype termed Multivulva (Muv).

Using the easily observed *C. elegans* Multivulva phenotype as a readout of Raf activity, we looked for genetic interactions between the DTS and weakly activating mutations in the N-terminal regulatory domains. By expressing mutant alleles in the VPCs we found that loss of the DTS enhanced LIN-45(S312A), a hyperactive mutant modeled after human *RAF1* mutations found in patients with Noonan Syndrome. We used this cellular context to perform a structure-function analysis and discovered three regions of the DTS required for normal regulation. In a complementary approach, we employed AlphaFold to predict the quaternary structure of LIN-45:14-3-3 complexes with regulatory phosphorylations and compared them with predictions of BRAF complexes from fly and human. The LIN-45 DTS is predicted to adopt a compact, T-shaped wedge between the kinase domain and the 14-3-3 protein while the fly and human DTS is predicted to adopt an extended conformation that makes extensive contacts with the kinase domain. The divergence between worm and the fly and human Raf DTS structures may represent distinct biophysical solutions to phosphorylation-gated negative feedback.

## Results

### Deletion of the LIN-45 DTS enhances LIN-45(S312A) activity

In human Raf proteins, mutations that prevent phosphorylation at the CR2 site disrupt 14-3-3 interaction, increase kinase activity, and cause disease. For example, mutations at or near the phosphorylated S259 residue of RAF1 are associated with NS (HEBRON *et al*. 2022). To facilitate screening for negative regulators of *C. elegans* LIN-45, we produced strains carrying single-copy transgenes that express YFP-tagged LIN-45(S312A), a mutation that prevents phosphorylation at the residue equivalent to RAF1 S259. All transgenes made in *C. elegans* were constructed using a *lin-31* promoter that drives expression in the VPCs during the L2 and L3 larval stages (MYERS AND GREENWALD 2005; TOWNLEY *et al*. 2023). Two independent strains carrying wild-type *yfp-lin-45(+)* had normal development, with no adults displaying the Muv phenotype (TOWNLEY *et al*. 2023) (Fig. 2a,b). All *yfp-lin-45(S312A)* strains were consistent, with a very small proportion of adults expressing the Muv phenotype, at a frequency of 0%, 0.6%, 1%, and 2% for four independent strains (Fig. 2a,b). Thus, the *yfp-lin-45(S312A)* transgene is well suited for genetic enhancement studies.

**Figure 2.**
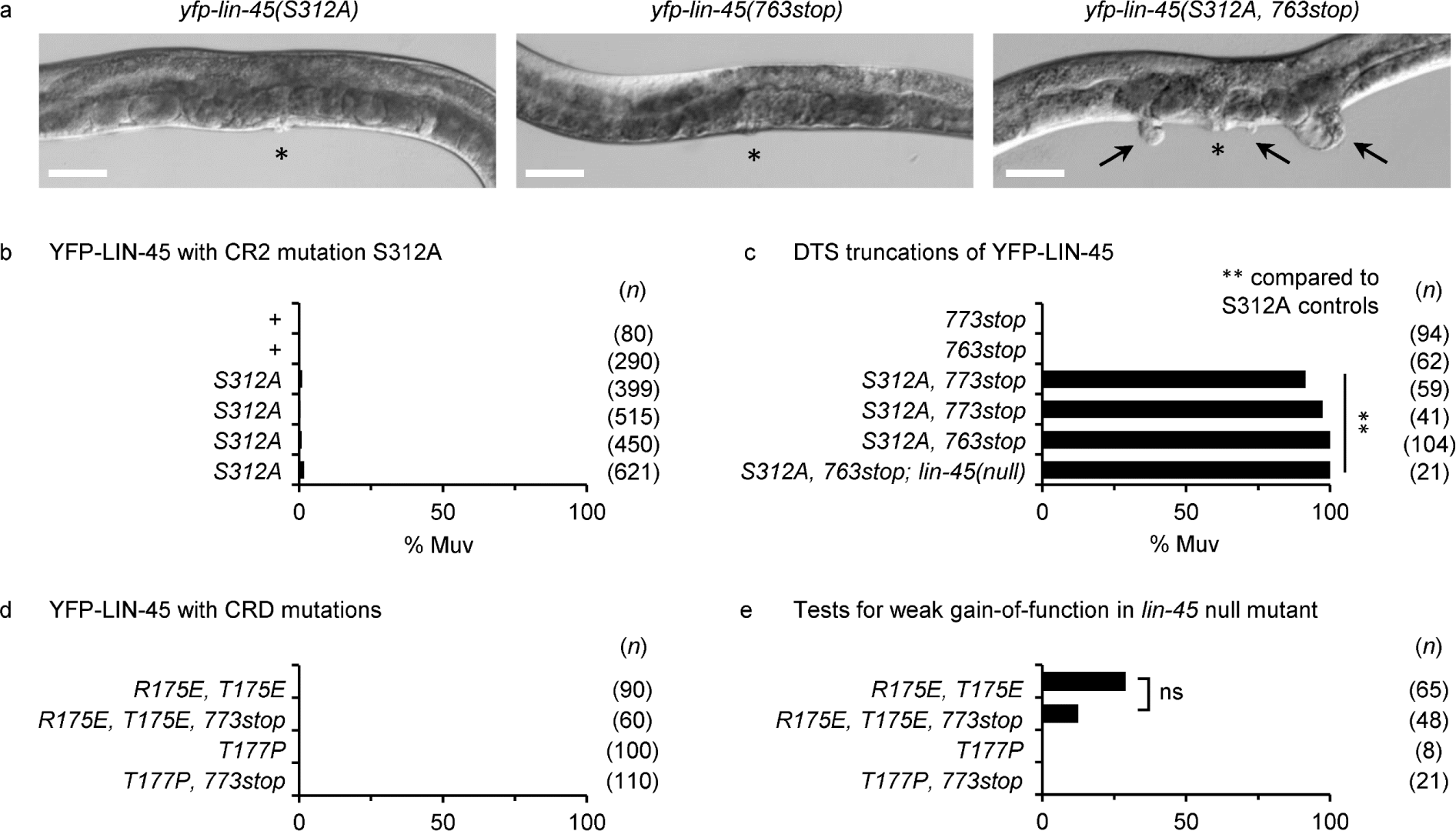
Deletion of the LIN-45 DTS enhances LIN-45(S312A) activity. **a)** Adult animals carrying transgenes expressing *yfp-lin-45(S312A)* (left), *yfp-lin-45(763stop)* (middle), or *yfp-lin-45(S312A, 763stop)* (right). Asterisks indicate the normal vulva; arrows indicate ectopic pseudovulvae. **b-e)** All panels are the percentage of adults displaying the Muv phenotype. Different, independent transgenic strains are listed as replicate genotypes. The number of adults scored (*n*) is shown in parentheses. **b)** Full-length LIN-45 transgenes *yfp-lin-45(+)* and *yfp-lin-45(S312A).* **c)** LIN-45 DTS truncations *yfp-lin-45(773stop), yfp-lin-45(763stop), yfp-lin-45(S312A, 773stop),* and *yfp-lin-45(S312A, 763stop).* The transgene *yfp-lin-45(S312A, 763stop)* was tested in the mutant *lin-45(dx19)*. Groups expressing YFP-LIN-45 with truncations were compared to YFP-LIN-45(S312A) controls using the Fisher’s exact test. **p-value <0.001. **d)** LIN-45 transgenes with CRD mutations *yfp-lin-45(R175E, T177E), yfp-lin-45(R175E, T177E, 773stop), yfp-lin-45(T177P),* and *yfp-lin-45(T177P, 773stop).* **e)** Weak gain-of-function activity was tested in the *lin-45(dx19)* mutant for *yfp-lin-45(R175E, T177E), yfp-lin-45(R175E, T177E, 773stop), yfp-lin-45(T177P),* and *yfp-lin-45(T177P, 773stop).* Those expressing YFP-LIN-45(R175E, T177E, 773stop) were compared to YFP-LIN-45(R175E, T177E) controls using the Fisher’s exact test. ns; not significant.

To test the role of the DTS in regulating LIN-45 activity, we generated new transgenes with C-terminal truncations, *yfp-lin-45(763stop)* and *yfp-lin-45(773stop),* endpoints which remove the DTS but retain the LIN-45 kinase domain and C-terminal 14-3-3 binding site. Neither truncated form produced phenotype defects (Fig. 2a,c). To test whether the DTS truncations enhance LIN-45(S312A), we produced *yfp-lin-45(S312A, 763stop)* and *yfp-lin-45(S312A, 773stop)* transgenes. Expression of either truncated mutant resulted in a highly penetrant Muv phenotype (100% of adults) (Fig. 2a,c). To test whether the LIN-45(S312A, L763stop) mutant can support signaling without wild-type *lin-45(+),* we introduced *yfp-lin-45(S312A, L763stop)* into the null mutant *lin-45(dx19).* When homozygous, this mutant is Vul, but can be completely rescued by *yfp-lin-45(+)* transgenes expressed in the VPCs (TOWNLEY *et al*. 2023). In *lin-45(dx19*) mutants, we observed a highly penetrant Muv phenotype in animals expressing *yfp-lin-45(S312A, L763stop)* (Fig. 2c). Together, these results indicate that the DTS is dispensable for kinase activation but important for negative regulation.

### Deletion of the LIN-45 DTS does not enhance mutations in the CRD

Structural studies indicate that the CR2 and CRD both interact with 14-3-3 and contribute to the inactive conformation (PARK *et al*. 2019). Mutations at two sites within the RAF1 CRD, R143E and T145E, disrupt its interaction with 14-3-3 (CLARK *et al*. 1997). We previously showed that mutations in the LIN-45 CRD, in transgenes expressing *yfp-lin-45(R175E, T177E),* cause weak gain-of-function activity in the *lin-45(dx19)* null mutant background (TOWNLEY *et al*. 2023). In BRAF, a single missense change at the same site, *BRAF(T241P),* is associated with CFC (SCHULZ *et al*. 2008).

We generated YFP-LIN-45 transgenes carrying either of two CRD mutations, a 14-3-3 interaction-disrupting double mutant R175E, T177E, or a single missense equivalent to that found in CFC, T177P. In a *lin-45(+)* genotype, none of the transgenes with CRD mutations caused defects, and the same was true of transgenes with a DTS truncation in *yfp-lin-45(R175E, T177E, 773Stop)* or *yfp-lin-45(T177P, 773Stop)* (Fig. 2d). To observe weak gain-of-function effects, we assayed each transgene in the homozygous *lin-45(dx19)* mutant. In the *lin-45(dx19)* mutant, a proportion of *yfp-lin-45(R175E, T177E, 773Stop)* animals displayed the Muv phenotype (13%); however, it was not significantly different from the proportion resulting from the *yfp-lin-45(R175E, T177E)* transgene in the same background (Fig. 1e). In the *lin-45(dx19)* mutant, no Muv phenotypes were observed in animals carrying the *yfp-lin-45(T177P, 773Stop)* transgene (Fig. 1e). We conclude that there is no genetic interaction between the *R175E, T177E* or *T177P* CRD mutations and loss of the LIN-45 DTS.

### Negative acting regulatory sequences within the LIN-45 DTS

To guide our analysis of regions within the LIN-45 DTS, we aligned a set of sequences that represent Raf proteins of nematodes, marine invertebrates, insects and vertebrates (Fig. 3a, Fig. S1). Compared to Raf proteins from other animals, the LIN-45 DTS contains two non-conserved insertions predicted to form helices (Fig. 3a,b). Despite this difference, we found similarities at two partially overlapping regions that were previously described in BRAF: the active site binding sequence (ASBS) and an ERK phosphorylation site we refer to as the “KTP motif.” We also found a third region of similarity at the extreme C-terminus, characterized by a motif consisting of aromatic residues, typically [FY]-x-x-[FYLI]. Using single-copy transgenes and the *lin-31* promoter strategy described above, we scanned the DTS region of *yfp-lin-45(S312A)*, introducing stop codons to test the requirement for each of these three sequence elements.

**Figure 3.**
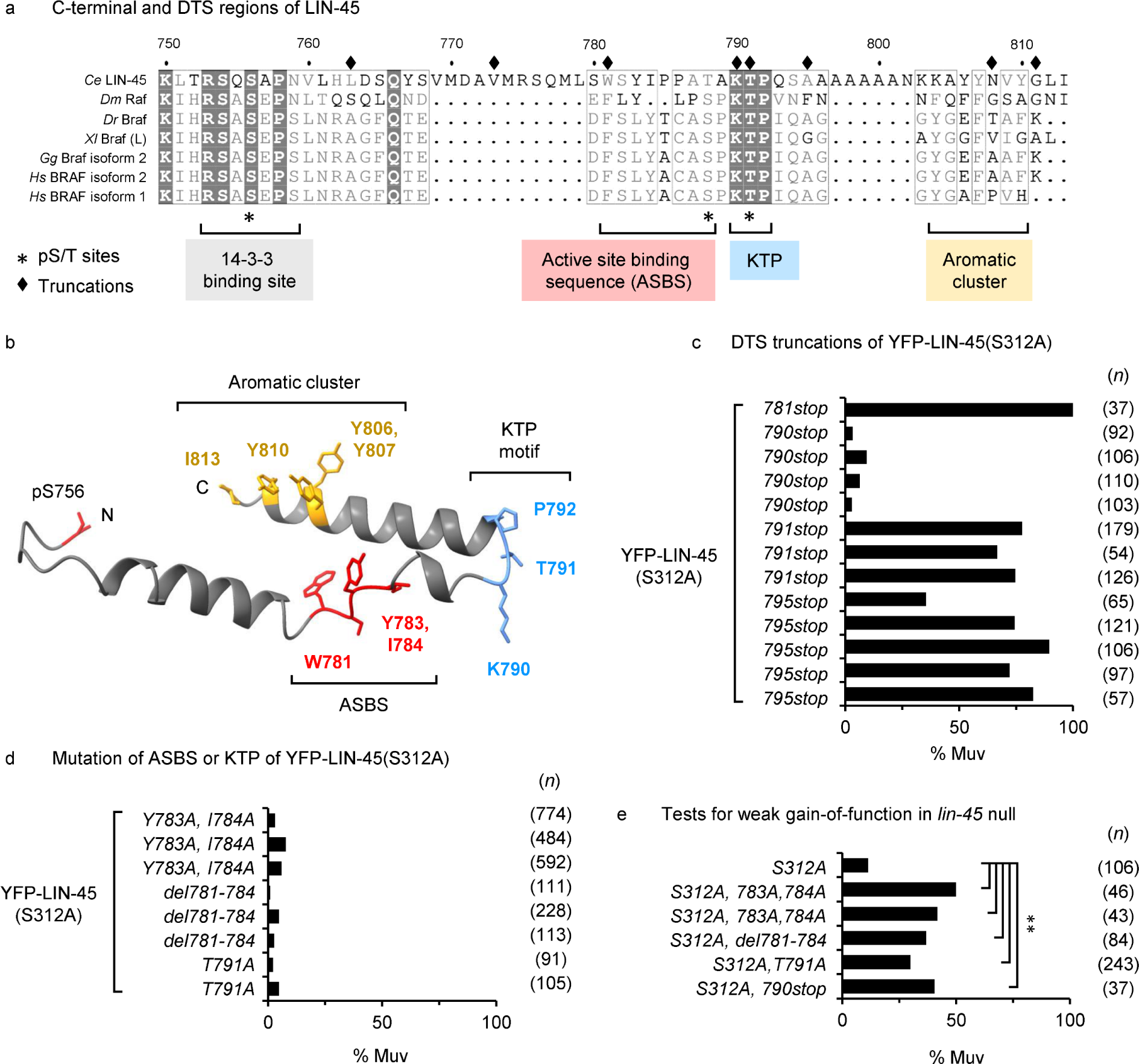
Negative-acting regulatory sequences within the LIN-45 DTS. **a)** Aligned C-terminal sequences of *C. elegans* LIN-45, *D. melanogaster* 621, *D. rerio* 733, *X. laevis* 757, *G. gallus* 765, *H. sapiens* isoforms 1 and 2 723. Numbering above alignment refers to LIN-45 protein. Asterisks indicate phospho-sites in human BRAF. Diamonds indicate stop codons introduced to truncate LIN-45. **b)** Predicted structure of LIN-45 region 756-813, from Alphafold model Q07292. R group sidechains are shown for residues of the ASBS (red), KTP motif (blue), and aromatic cluster (yellow). **c-e)** All panels are the percentage of adults displaying the Muv phenotype. Different, independent transgenic strains are listed as replicate genotypes. The number of adults scored (*n*) is shown in parentheses. **c)** DTS truncations in LIN-45(S312A) transgenes *yfp-lin-45(S312A, 781stop), yfp-lin-45(S312A, 790stop), yfp-lin-45(S312A, 791stop)*, and *yfp-lin-45(S312A, 795stop).* **d)** Mutations in LIN-45(S312A) transgenes *yfp-lin-45(S312A, Y783A, I784A), yfp-lin-45(S312A, del781-784), yfp-lin-45(S312A, T791A).* **e)** Weak gain-of-function activity was tested in the *lin-45(dx19)* mutant for *yfp-lin-45(S312A), yfp-lin-45(S312A, Y783A, I784A), yfp-lin-45(S312A, del781-784),* and *yfp-lin-45(S312A, T791A).* All groups were compared to those expressing YFP-LIN-45(S312A) using the Fisher’s exact test. **p-value <0.001.

### Tests of the active site binding sequence

In cryo-EM structures of BRAF complexed with 14-3-3 proteins, the BRAF DTS residues 743-750 (FSLYACAS) interact in trans to occupy the ATP-binding pocket of the second BRAF protomer (KONDO *et al*. 2019). While not a strictly conserved sequence, alignment of the ASBS of vertebrate and invertebrate Raf proteins revealed this region tends to have one or more aromatic, hydrophobic, and bulky residues, with the most conserved position at BRAF F743 (Fig. S2a). We term this position of the ASBS “1.” LIN-45 has two aromatic residues in this interval (781-788, WSYIPPAT): W781 at position 1 and Y783 at position 3 (Fig. 3a, Fig. S2a).

To remove all LIN-45 DTS elements including the ASBS, we introduced a stop codon at position 781. Like the DTS truncations at 763 and 773, a transgene expressing *yfp-lin-45(S312A, 781stop)* caused a highly penetrant Muv phenotype, with nearly 100% of animals displaying the phenotype (Fig. 3c). To test a truncation that retains the ASBS but lacks the KTP motif and C-terminal aromatic cluster, we introduced a stop codon at position 790. Animals carrying *yfp-lin-45(S312A, 790stop)* transgenes displayed a low penetrance of the Muv phenotype, at a frequency of 3%, 3%, 4%, and 9% for four independent strains (Fig. 3c). To confirm that YFP-LIN-45(S312A, 790stop) protein is expressed in these strains, we measured YFP fluorescence intensity in VPCs at the L2 larval stage for animals carrying *yfp-lin-45(S312A, 790stop)*, as well as other transgenes that caused either low or highly penetrant Muv phenotypes (Fig. S3a). We did not observe a correlation between measured YFP fluorescence intensity and the penetrance of the Muv phenotype (Fig. S3b). Thus, the failure of *yfp-lin-45(S312A, 790stop)* to cause the Muv phenotype results from its intrinsic activity rather than expression level. These results indicate that residues 781-789 of LIN-45 contain negative regulatory activity.

### The active site binding sequence contributes to negative regulation

To guide our mutagenesis of the LIN-45 ASBS element, we superimposed the cryo-EM model of the BRAF DTS (KONDO *et al*. 2019) and the x-ray diffraction model of BRAF bound to an ATP analog AMP-PCP (LIAU *et al*. 2020b) (Fig. S2a). BRAF binds the adenine rings of ATP at a hydrophobic pocket in the kinase domain (HANKS AND HUNTER 1995; TSAI *et al*. 2008). In the cryo-EM model, the same pocket is occupied by the BRAF DTS peptide sequence FSLYACA (Fig. S2b). Superposition of the models suggested that BRAF DTS residues L745 and Y746 occupy the same location as ATP, and that the same positions in LIN-45, Y783 and I784, were likely candidates for function-disrupting mutagenesis (Fig. S3a,b).

We engineered two sets of mutations to test the putative ASBS in LIN-45. In the first transgene, *yfp-lin-45(S312A, Y783A, I784A)*, residues at positions 2 and 3 were mutated to alanine. In the second, *yfp-lin-45(S312A, del W781-I784)*, we deleted four predicted ASBS residues, 781-784. The proportion of Muv animals in strains carrying *yfp-lin-45(S312A, Y783A, I784A)* transgenes was low, at a frequency of 3.0%, 5.9%, and 7.9% in independent strains (Fig. 3d). Strains carrying the deletion *yfp-lin-45(S312A, del W781-I784)* strains also had a low penetrance of the Muv phenotype, at a frequency of 0.9%, 2.7%, and 4.8% in independent strains (Fig. 3d). We next tested the possibility that these mutants are weakly gain-of-function by introducing them into the *lin-45(dx19)* null mutant. Whereas 11% of animals carrying the control *yfp-lin-45(S312A)* transgene in *lin-45(dx19)* mutants were Muv, two independent *yfp-lin-45(S312A, Y783A, I784A)* transgenes resulted in significantly higher penetrances, at 50% and 42%, in the *lin-45(dx19)* mutants (Fig. 3e). Animals carrying the deletion transgene *yfp-lin-45(S312A, del W781-I784)* in the *lin-45(dx19)* mutant also displayed a significantly increased penetrance at 37% of animals (Fig. 3e). From these results we conclude that LIN-45 ASBS residues, specifically Y783 and I784, act to negatively regulate signaling.

### Tests of the KTP feedback phosphorylation site

To investigate the role of the LIN-45 KTP motif, we made *yfp-lin-45(S312A)* transgenes truncated at position 795, which retains the ASBS and KTP motif but removes the C-terminal aromatic cluster. Because phosphorylation of the BRAF KTP motif is inhibitory, we expected that *yfp-lin-45(S312A, 795stop)* transgenes would be similar to or less active than the truncation at K790. However, animals with this allele displayed a highly penetrant Muv phenotype, at a frequency of 35%, 72%, 74%, 82%, 90% in five independent strains (Fig. 3b). The LIN-45 KTP motif is centered at residue T791, equivalent to the ERK phosphorylation site T753 in BRAF. To determine whether the presence of this threonine was responsible for the surprising behavior of *yfp-lin-45(S312A, 795stop)*, we tested *yfp-lin-45(S312A, 791stop)*, a truncation that retained K790 but removed T791. Strains expressing this allele also exhibited a Muv phenotype, at a frequency of 67% and 75% in independent strains (Fig. 3b). This observation was unexpected, and in principle could suggest that the KTP motif enhances activity.

### The KTP motif residue T791 is required for negative regulation

To more specifically test the role of the KTP motif, we generated the transgene *yfp-lin-45(S312A, T791A)*, a mutation that prevents phosphorylation of T791. The two isolated strains displayed the Muv phenotype at low penetrance, at a frequency of 2% and 5% (Fig. 3d). To better assess its gain-of-function activity, we introduced the weakest of these transgenes into the *lin-45(dx19)* mutant, where the penetrance of the Muv phenotype was 30%, a significant increase compared to the *yfp-lin-45(S312A); lin-45(dx19)* control (Fig. 3e). We also tested the transgene *yfp*-*lin-45(S312A, 790stop)* because this truncation lacks the KTP motif. The penetrance of the Muv phenotype resulting from *yfp*-*lin-45(S312A, 790stop)* in *lin-45(dx19)* was significantly enhanced, at a frequency of 41% (Fig. 3e). These results support the interpretation that T791 and potentially other sequences still C-terminal to the ASBS negatively regulate LIN-45 signaling.

Because missense mutations disrupting either the ASBS or KTP motif in the context of full-length LIN-45(S312A) resulted in increased activity, we concluded that both regions act in negative regulation. It is striking that individual mutations in the ASBS or KTP motif caused weak gain-of-function, in contrast to the strong activity of almost all truncations we tested. This suggests that sequences still more C-terminal to position 795 make a large contribution to negative regulation. The notable exception to this trend was a truncation at the end of the ASBS region, position 790.

### Tests of the C-terminal aromatic cluster

We investigated the role of a cluster of aromatic residues at the extreme C-terminus of LIN-45, residues 806-813 (YYNVYGLI). To remove the most C-terminal tyrosine of the aromatic cluster, we introduced a stop codon at N808. Expression of *yfp-lin-45(S312A, 808stop)* resulted in a highly penetrant Muv phenotype, at a frequency of 55%, 86%, and 87% in three independent strains (Fig. 4a). We also tested a form that terminates after the tyrosines, by introducing a stop codon at G811. Expression of *yfp-lin-45(S312A, 811stop)* caused a Muv phenotype, at a frequency of 46%, 85%, 90%, and 92% in four independent strains (Fig. 4a). These results suggest that sequences within the C-terminal interval 808-813 are necessary for negative regulation of LIN-45(S312A).

**Figure 4.**
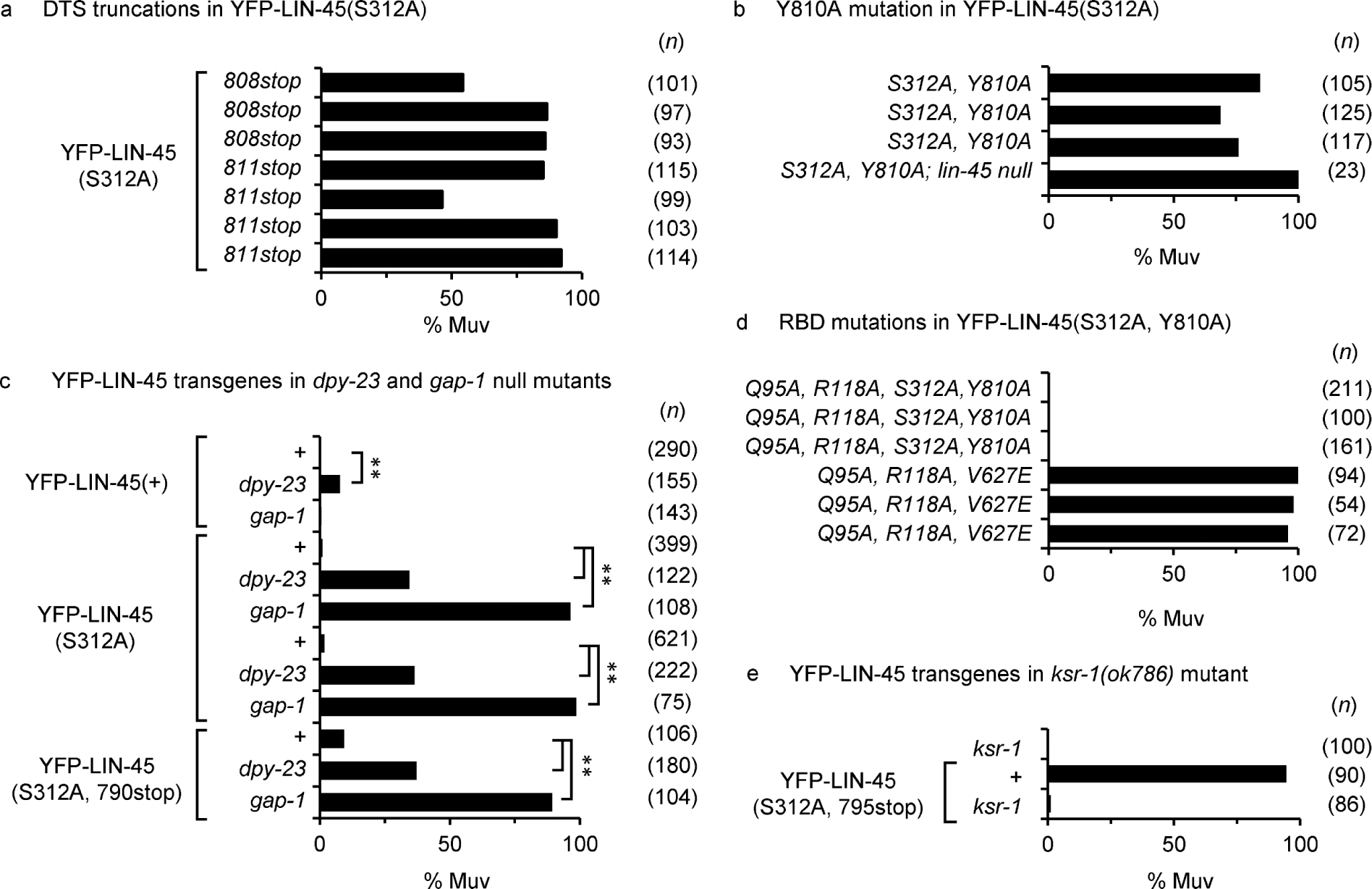
A C-terminal aromatic cluster is critical for LIN-45 negative regulation. All panels are the percentage of adults displaying the Muv phenotype. Different, independent transgenic strains are listed as replicate genotypes. The number of adults scored (*n*) is shown in parentheses. **a)** DTS truncations in LIN-45(S312A) transgenes *yfp-lin-45(S312A, 808stop), yfp-lin-45(S312A, 811stop).* **b)** Mutations in LIN-45(S312A) transgenes *yfp-lin-45(S312A, Y810A).* The transgene *yfp-lin-45(S312A, Y810A)* was tested in the mutant *lin-45(dx19)*. **c)** Tests of *dpy-23*/AP2M1 and *gap-1*/GAP mutants. Transgenes *yfp-lin-45(+), yfp-lin-45(S312A), yfp-lin-45(S312A, 790stop)* were introduced into *dpy-23(e840)* and *gap-1(ga133)* mutants. For all groups expressing the same transgene, *dpy-23(e840),* and *gap-1(ga133)* mutants were compared to wild-type controls using the Fisher’s exact test. **p-value <0.0001. d) Mutation of the RBD was tested in *yfp-lin-45(Q95A,R119A,S312A,Y810A).* e) The transgene *yfp-lin-45(+), yfp-lin-45(S312A, 795stop)* was introduced into *ksr-1(ok786)* mutants.

### The aromatic cluster sequence residue Y810 is critical for negative regulation

In LIN-45, Y810 is located in the aromatic cluster and at the nexus of two motifs, a Y-x-x-Y motif spanning 807-810, and a Y-G-x-[YFLI] motif at 810-813 (Fig. 2a). When present at the C-terminus of protein substrates, both motifs are associated with endocytosis, with the Y-G-x-[YFLI] motif specific to interaction with the AP-2 subunit mu protein AP2M1. To test whether Y810 is necessary for LIN-45 negative regulation, we generated the *yfp-lin-45(S312A, Y810A)* transgenes. Animals harboring *yfp-lin-45(S312A, Y810A)* transgenes expressed the Muv phenotype with a penetrance that varied from 69%, 87% and 84% in independent strains (Fig. 4b). The penetrance of the Muv phenotype increased to 100% when this transgene was assessed in the *lin-45(dx19)* null mutant. These results indicate that Y810 is critical for negative regulation.

Endocytosis has not been described as regulating Raf activity, although Ras (ROY *et al*. 2002), EGFR (CARPENTER 2000), RAF1 and MEK (POL *et al*. 1998) are endocytosed during signaling. We reasoned that if the C-terminal aromatic cluster acts as an internalization signal, then loss the *C. elegans* AP2M1 ortholog *dpy-23* may enhance the Muv phenotype of *yfp-lin-45(S312A).* We combined the null allele *dpy-23(e840)* with *yfp-lin-45(S312A)* or wild-type *yfp-lin-45(+)* transgenes. Genotypes with *dpy-23(e840); yfp-lin-45(S312A)* displayed Muv phenotypes in 21% and 27% of animals, a significant increase from either of the two *yfp-lin-45(S312A)* transgenes alone (Fig. 4c). However, genotypes carrying *dpy-23(e840); yfp-lin-45(+)* also displayed Muv phenotypes in 12% of animals, a significant enhancement compared to the strain carrying *yfp-lin-45* alone (Fig. 4c). Thus, *dpy-23(e840)* enhances both *yfp-lin-45(S312A)* and wild-type *yfp-lin-45(+)*. Because DPY-23/AP2M1 promotes internalization of many substrates, we were concerned about pleiotropic effects of this mutant. To assess whether loss of *dpy-23* has a direct effect on YFP-LIN-45(S312A) through the DTS region, we tested the effect of *dpy-23(e840)* on *yfp-lin-45(S312A, 790stop)* transgene, an allele lacking all putative C-terminal internalization signals. In the *dpy-23(e840)*; *yfp-lin-45(S312A, 790stop)* genotype, we observed 26% of animals displayed Muv phenotypes, a significant difference from *yfp-lin-45(S312A, 790stop)* alone (Fig. 4c). These results indicate that *dpy-23* loss enhances LIN-45 signaling activity; however, they do not support a role for the DTS and aromatic cluster in internalization.

### LIN-45(S312A) signaling is dependent on Ras and KSR activity

Dephosphorylation of the CR2 region of Raf is thought to disrupt 14-3-3 interaction, destabilize the autoinhibited conformation, and promote kinase dimerization (SENDOH *et al*. 2000; RITT *et al*. 2010). To better understand how mutations in the DTS increase LIN-45(S312A) activity, we tested for Ras dependence by introducing mutations in the LIN-45 RBD that prevent Ras binding. Residues Q95 and R118 of the LIN-45 RBD correspond to Q66 and R89 of RAF1, sites that are required for interaction with Ras-GTP and plasma membrane recruitment (BLOCK *et al*. 1996; HSU *et al*. 2002; HARDING *et al*. 2003). We previously showed that transgenes expressing YFP-LIN-45(Q95A, R118A) fail to rescue the *lin-45* null mutant, consistent with a loss of function (TOWNLEY *et al*. 2023). To test whether a highly active LIN-45(S312A) form requires Ras binding, we generated the transgene *yfp-lin-45(Q95A, R118A, S312A, Y810A)*. Animals carrying this transgene did not exhibit the Muv phenotype in three independent strains (Fig. 4d), indicating that the S312A and Y810A mutations do not bypass Ras-GTP requirement. This is not a trivial result, as the Muv phenotype caused by LIN-45(V627E) mutant, which carries an activation loop mutation, was not suppressed by introduction of Q95A and R118A (Fig. 4d). The result that LIN-45(S312A, Y810A) form is Ras dependent is consistent with a role for Ras-GTP in mediating dimerization and activation.

We also investigated whether the LIN-45(S312A) mutant is affected by elevated Ras-GTP levels. The gene *gap-1* encodes a GTPase Activating Protein (GAP). While loss of *gap-1* alone does not cause Muv phenotypes, it is known to enhance other mutants that compromise negative regulation of Raf or ERK signaling (YOO *et al*. 2004; DE LA COVA AND GREENWALD 2012). We introduced *yfp-lin-45(S312A)* and *yfp-lin-45(+)* transgenes into the null mutant *gap-1(ga133),* generating *yfp-lin-45(S312A); gap-1(ga133)* and *yfp-lin-45(+); gap-1(ga133)*. Each of the *yfp-lin-45(S312A); gap-1(ga133)* strains had a highly penetrant Muv phenotype, at a frequency of 96% and 99%, while the *yfp-lin-45(+); gap-1(ga133)* genotype displayed completely normal development (Fig.4c). Taken together, the results of mutations in the LIN-45 RBD and the *gap-1* mutant indicate that the LIN-45(S312A) form is dependent and highly sensitive to Ras-GTP.

Kinase Suppressor of Ras (KSR) is a conserved binding partner of Raf that is required for MEK phosphorylation (LAVOIE *et al*. 2018). Once Raf is dephosphorylated at the CR2 site, KSR and Raf can form heterodimers, which appears to promote Raf activation through its dissociation from an inhibitory interaction with MEK (BONED DEL RIO *et al*. 2019). Genetically, loss of *ksr-1* is epistatic to an activated form of LIN-45 that has mutations in the activation loop (ROCHELEAU *et al*. 2002). We expected that the highly active LIN-45 mutants we tested would require KSR-1 activity to cause the Multivulva phenotype. Indeed, when we combined the null allele *ksr-1(ok786)* with *yfp-lin-45(S312A, 795stop)*, the highly penetrant Muv phenotype caused by this allele was completely suppressed (Fig. 4e). These results indicate that LIN-45 lacking the DTS is activated by normal KSR-dependent steps.

### Predicted quaternary structure of LIN-45 DTS, kinase domain, and 14-3-3

We used Alphafold 3 to predict the quaternary structure of heterotetramers containing LIN-45 and PAR-5, an ortholog of 14-3-3 zeta. Because Alphafold permits introduction of post-translational modifications (ABRAMSON *et al*. 2024), we used it to predict the consequences of T791 phosphorylation and DTS mutations. Alphafold generates an “ensemble” of five models and a residue-by-residue measure of the confidence for the structure. We used the following criteria for assessing the predicted models: i) agreement between the five ensemble models, ii) the pLDDT confidence score, iii) how well the model responded to phosphorylation or genetic changes that we tested experimentally.

We queried Alphafold with two copies of LIN-45, residues 470-813 and phosphorylated at the C-terminal 14-3-3 binding sequence, and two copies of PAR-5. The query returned a quaternary structure in good agreement with multiple, experimentally derived cryo-EM structures that show a symmetric heterotetramer of BRAF and 14-3-3 proteins (PARK *et al*. 2019). The dimer of PAR-5 adopts a two-sided cradle with a symmetrical dimer of LIN-45 kinase domains tethered at pS756 (Fig.5a,b, Fig.S5a). Prediction confidence within much of the LIN-45 kinase domain and PAR-5 was high, with pLDDT scores >90. However, prediction confidence for the DTS region was very low, with pLDDT scores <50, and the five models of the DTS did not overlap well in their predicted location (Fig. S5b, Fig.S6a).

We next introduced phosphorylations in LIN-45, at T626 and T629 of the activation loop, and T791 of the DTS. With all phosphorylations present, we observed increased agreement between the five ensemble models with regard to DTS location and orientation, and increased pLDDT scores (Fig.5a,b, Fig.S6a-d). In this form, the DTS adopted a compact, T-shaped structure in which the helices interact with one another and occupy the space between PAR-5 and the kinase domain ATP-binding pocket (Fig. 5b). In all five ensemble models, residues of the aromatic cluster, including Y810, occupied a complimentary pocket on PAR-5, the KTP motif formed a sharp turn, and the ASBS residues 778-783 interacted with the kinase domain near the ATP-binding pocket (Fig. 5b). Predictions of LIN-45 with activation loop phosphorylation but no modification of T791 produced five models that differed from one another with regard to DTS location and orientation (Fig. 5a). Comparison of these two predictions suggests that DTS phosphorylation acts as a switch, causing the DTS to adopt a compact conformation and interact with the kinase domain (Fig. 5a,b).

**Figure 5.**
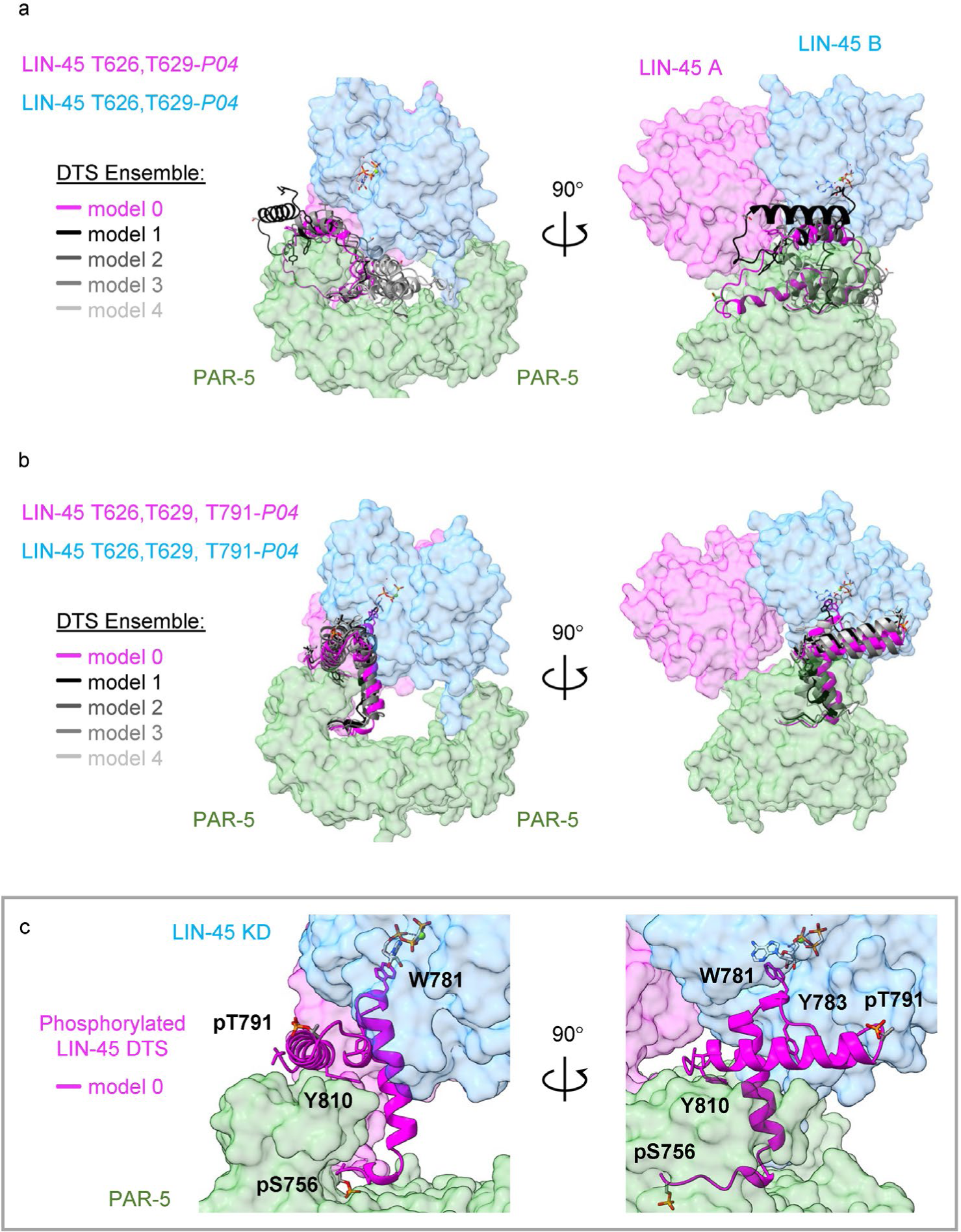
Predicted trans-interaction of the LIN-45 DTS and kinase domain. All panels show two orthogonal views of models predicted by AlphaFold for the heterotetramer containing two protomers of LIN-45 residues 470-813 with pS756 (protomers A and B in magenta and blue), and two protomers of PAR-5 (green). For the LIN-45 DTS, panels a-b show five ensemble models superimposed and depicted as ribbons with the highest confidence model in magenta. For clarity, PAR-5 is trimmed to exclude the C-terminal tail (residues 230-248). To indicate the location of the ATP binding pocket, ATP is superimposed on the kinase domain models. a) LIN-45 with phosphorylation at the activation loop (pT626, pT629) and unphosphorylated DTS. b) LIN-45 with phosphorylation at the activation (pT626, pT629) and the DTS (pT791). c) Higher magnification views of model shown in panel b, with only the highest confidence model depicted in ribbon (magenta). The locations of residues in the ASBS (W781), KTP motif (pT791), and aromatic cluster (Y810) are indicated.

We reasoned that AlphaFold may predict structural changes that correspond to the weak and strong phenotypes resulting from mutations we tested *in vivo*. We examined the effects of weak gain-of-function alleles, T791, del781-784, and Y783A, I784A. Models of LIN-45 T791A resembled those of wild-type without phosphorylation, meaning the DTS failed to form a compact shape, did not interact with the kinase domain, and there was little agreement between the models (Fig. S7a,b). Similarly, models of LIN-45 del781-784 showed poor agreement and failed to adopt a compact conformation (Fig. S7d). Models of LIN-45 Y783A, I784A were well ordered and in strong agreement with one another. However, the ASBS region adopted an α-helical conformation and rotated 90 degrees away from the kinase active site (Fig.S7c).

We next tested the strong gain-of-function alleles Y810A and 811stop, which affect the aromatic cluster. LIN-45 Y810A models were not in agreement with one another nor did the DTS adopt a compact structure (Fig. S7e). LIN-45 811stop models showed some degree of agreement and the DTS adopted secondary structure, but the phospho-KTP turn was rotated 180 degrees compared to wild type (Fig. S7f). Comparison of these predictions to each other and wild type suggests an important role for Y810 in stabilizing the secondary structure of the DTS and a role for the C-terminal sequence at G811 (GLI) in anchoring the DTS to PAR-5.

Lastly, we used AlphaFold to understand how the truncation at 790 produced weak gain-of-function while a truncation at 791 caused strong gain-of-function. Predictions of LIN-45(790stop) produced two out of five models with the kinase active site occupied by the ASBS region (Fig. S7g). By contrast, all models of LIN-45(791stop) were in strong agreement, with the DTS oriented away from the kinase domain (Fig. S7h). Overall, AlphaFold predictions using alleles that caused phenotypic consequences *in vivo* tended to disrupt the protein-protein interactions between the DTS and either the kinase domain or 14-3-3.

### Predictions for human and fly Raf proteins

We queried AlphaFold with sequences of human BRAF isoform 1, BRAF isoform 2, or *Drosophila* Raf. The alternate C-termini of BRAF isoform 1 and 2 are conserved across vertebrates, with BRAF isoform 1 terminating in Y-G-A-F-P-V-H while isoform 2 terminates in Y-G-E-F-A-A-F-K (Fig. 3a, Fig. S1). *Drosophila* Raf also has a C-terminal interval containing multiple aromatic residues (Fig. S1). For all three of these Raf proteins, AlphaFold predicted the kinase domains to be in a symmetrical dimer within a dimer of 14-3-3 proteins (Fig. 6a-c). In the human BRAF isoform 1 predictions, the ensemble of DTS models were in poor agreement and none showed interaction of the ASBS residues with the kinase domain (Fig. 6a). Some models of BRAF isoform 1 predicted interactions between the phosphorylated residues pS750 and pT753 and the ATP binding pocket, and between the aromatic cluster residues Y760 and F763 and the kinase α-G helix, a region that binds MEK (Haling, 2014). By contrast, models of BRAF isoform 2 were in strong agreement with one other (Fig. 6b). Furthermore, the BRAF isoform 2 ASBS residues 743-746 (FSLY) were predicted to occupy the ATP binding pocket in strong agreement with the cryo-EM structure (KONDO *et al*. 2019), the phosphorylated residues pS750 and pT753 interacted with the surface of the kinase domain and the aromatic cluster interacted with the pocket formed by the α-G helix (Fig. 6b). Models of *Drosophila* Raf strongly agreed with one another and were very similar to those predicted for human BRAF isoform 2 (Fig. 6c). Although models for human and *Drosophila* Raf showed considerable differences compared to those for LIN-45, it was notable that the three DTS elements, the ASBS, the KTP motif, and the aromatic cluster, were predicted to participate in DTS-kinase domain interactions, suggesting some conservation of DTS functions.

**Figure 6.**
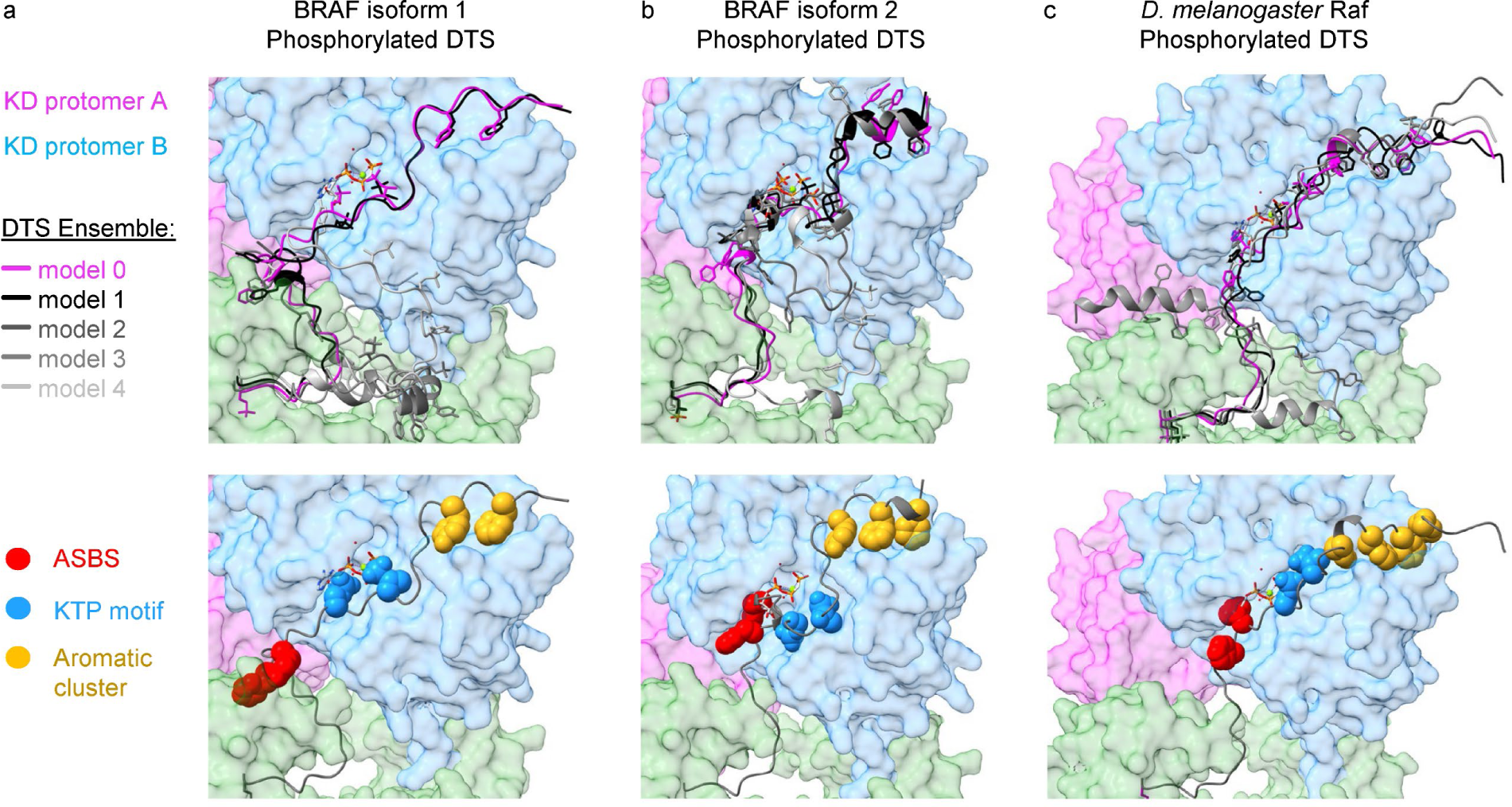
Comparison of the predicted trans-interactions for the DTS and kinase domain of human and fly Rafs. All panels show models predicted by AlphaFold for the heterotetramer containing two protomers of Raf kinase domain (protomers A and B in magenta and blue), and two protomers of 14-3-3 zeta (green). Top panels show five ensemble models superimposed and depicted as ribbons with the highest confidence model in magenta. Bottom panels show the residue sidechains in the ASBS (red), KTP motif (blue), and aromatic cluster (yellow) depicted in spheres. To indicate the location of the ATP binding pocket, ATP is superimposed on the kinase domain models. a) Human BRAF isoform 1 with phosphorylation at the DTS (pS750, pT753). b) Human BRAF isoform 2 with phosphorylation at the DTS (pS750, pT753). c) *D. melanogaster* Raf with phosphorylation at the DTS (pS666, pT669).

## Discussion

This work demonstrates that the *C. elegans* Raf/LIN-45 C-terminal DTS acts to negatively regulate Raf signaling. We showed that truncation of the DTS did not cause a phenotype alone but resulted in strong activation when combined with a mildly activating CR2 mutant, LIN-45(S312A). Experiments in mammalian cells previously established a negative regulatory role for phosphorylation of the Raf DTS by ERK (BRUMMER *et al*. 2003; RUSHWORTH *et al*. 2006; RITT *et al*. 2010). In this work, we tested nonsense and missense mutants of LIN-45 to further define the regulatory features within the DTS. These include three elements: the kinase domain-interacting sequences (ASBS), the KTP motif site of ERK phosphorylation, and the aromatic cluster.

Because of the wealth of structural and biochemical data on mammalian Rafs and our genetic analysis of the LIN-45 DTS mutant phenotypes, we leveraged structure prediction by AlphaFold to visualize activated heterotetramer complexes containing Raf and 14-3-3 proteins from worm, fly and human. Combining our predictions of LIN-45 structure with our experimental data set, we propose a feedback inhibition model for the function of the LIN-45 DTS. We propose that the ASBS acts as a kinase inhibitor that functions by binding the kinase domain. Phosphorylation of the KTP motif at T791 of LIN-45 acts to switch the DTS into the compact shape needed for inhibition. Finally, the aromatic cluster acts to anchor the inhibitory conformation by binding within the DTS and the 14-3-3 protein. In this discussion, we support these interpretations with insights gained from our analysis of LIN-45 mutants, and we propose that this model might represent a unique biophysical solution distinct from how mammalian and ancestral BRAF DTS functions.

### Role of the active site binding sequence (ASBS)

Our data provide evidence that the LIN-45 ASBS inhibits signaling. The strongest gain-of-function alleles we tested were DTS truncations that remove the ASBS along with all other DTS sequences from LIN-45(S312A). Shorter truncations that retained the ASBS were less active than deletion of the entire DTS. In the context of full-length LIN-45(S312A), missense mutations or a small deletion of the ASBS also caused gain-of-function, but with a weaker effect that was significant only when these mutants were assayed in the *lin-45* null background. Two previous studies tested a truncation in human BRAF that is equivalent to the LIN-45(763stop) mutation; neither observed a significant difference in signaling as measured by phospho-ERK levels (KONDO *et al*. 2019; LIAU *et al*. 2020a). Neither of these studies combined DTS deletion with a second gain-of-function mutation analogous to LIN-45(S312A) as we did in this study. We think it is likely that effects on BRAF signaling caused by truncation of the DTS will exhibit dependence on additional, weak gain-of-function mutations, as this was already shown to be true for mutation of the phospho-sites S750 and T753 (RITT *et al*. 2010).

The amino acid sequence of the ASBS is not strictly conserved but is enriched in aromatic, bulky, and hydrophobic residues (Fig. S1, Fig. S2). In most orthologs, including LIN-45, the ASBS contains two aromatic residues in a [FYW]-x-[FY] or [FYW]-x-x-[FY] motif. Structures of LIN-45 and PAR-5 predicted by AlphaFold position the LIN-45 ASBS residues matching this motif (at W781, sequence WSY) adjacent to the ATP-binding pocket of the kinase domain. These models predict that the LIN-45 DTS acts as an autoinhibitor and would suggest that a short motif of aromatic, bulky residues in other animal Raf DTS share a common function (Fig. S1, Fig. S2).

### Role of the KTP motif

Introduction of the T791A mutation to full-length LIN-45(S312A) caused weak gain-of-function that is apparent in the *lin-45* null background. The KTP motif is nearly invariant across animal Raf proteins, and the phospho-sites S750 and T753 of BRAF were already established as direct targets of ERK phosphorylation (RITT *et al*. 2010). In BRAF, mutation of S750 or T753 resulted in increased kinase activity and BRAF-RAF1 heterodimerization (BRUMMER *et al*. 2003; RUSHWORTH *et al*. 2006; RITT *et al*. 2010) This raises the question of what specific interactions are altered by phosphorylation. Based on the structure predictions for phosphorylated and un-phosphorylated LIN-45 complexes, we propose that phosphorylation at T791 allows the DTS to form a compact shape and interact with the kinase domain and PAR-5. In this model, pT791 is not a direct kinase domain binding site but rather a “switch” that promotes the DTS conformation change. Importantly, our mutation analysis indicated that the KTP motif alone is not sufficient for regulation; rather, the entire DTS must be present.

### Role of the aromatic cluster

We refer to LIN-45 residues 806-813 as the aromatic cluster because this interval contains multiple tyrosine residues. Of all the missense changes we tested in the LIN-45 DTS, mutation Y810A had the strongest phenotypic effect. This motif of aromatic residues is present in many other animals, as either [FY]-x-x-[FY] or [FY]-x-x-[FY]-x-x-[FYILMV] (Fig. S1, Fig. S3). A function for the aromatic cluster was not previously described. We first investigated whether the aromatic cluster acts as an internalization signal, finding that LIN-45(S312A) signaling was enhanced by the loss of *dpy-23*/AP2M1. However, enhancement was independent of the DTS, inconsistent with a function as an internalization signal. Prediction of the structure of complexes with LIN-45 and PAR-5 provided an alternative possibility. Y810 and other aromatic cluster residues were predicted to bridge the two helices of the DTS between PAR-5 and the kinase domain. We propose that these interactions are required as an “anchor” that may stabilize the inhibitory DTS structure in the space between the 14-3-3 side and the kinase domain.

### Comparative analysis of DTS structure predictions for worm, fly, and human

AlphaFold predicted different DTS conformations and subunit contacts for *C. elegans* LIN-45 compared to human BRAFs or *Drosophila* Raf. LIN-45 DTS structure responded to T791 phosphorylation by adopting a T-shaped conformation that occupied the space between the kinase domain and the 14-3-3 protein. By contrast, the DTS of human and fly Raf adopted an extended conformation with contacts along the entire kinase surface anchored by the C-terminal aromatic cluster to a pocket formed by the kinase α-G helix that functions as a docking point for MEK Interestingly, the DTS of worm, fly and human Rafs all appeared to utilize the ASBS to bind the kinase domain at sites near the ATP-binding pocket. However, the specific contacts predicted for the KTP motif and aromatic cluster differed between LIN-45 and other Rafs. For example, in LIN-45, residues of the aromatic cluster anchored the DTS C-terminus to PAR-5. In *Drosophila* Raf and human BRAF, aromatic cluster residues anchored the DTS C-terminus into a pocket on the kinase domain, a location where it could potentially block contact between BRAF and MEK. Our AlphaFold predictions also suggest some differences between the two BRAF isoforms, which was manifested in low model agreement for BRAF isoform 1 and high agreement for isoform 2. The most well-studied form, BRAF isoform 1, ends in a short aromatic sequence conforming to a [FY]-x-x-[FY] motif, while BRAF isoform 2 contains a longer motif with additional aromatic residues (Fig. 3a) (MARRANCI *et al*. 2017). The modeled differences in structures of BRAF isoform 1 and 2 suggest they may have distinct biochemical properties and responses to feedback inhibition, thus providing additional complexity to Raf signaling.

## Materials and Methods

### C. elegans genetics

The genotypes of *C. elegans* strains used in this work are listed in Table S1. The following alleles were used: LG IV: *lin-45(dx19).* LG X: *dpy-23(e840), gap-1(ga133), ksr-1(ok786)*. In strains carrying *lin-45(dx19),* this allele was maintained as a heterozygous genotype *lin-45(dx19)/nT1 IV; +/nT1[umnIs28] V* and homozygous *lin-45(dx19)* animals were identified by the absence of the GFP-marked balancer.

### Plasmids and transgenes for YFP-LIN-45 expression in *C. elegans*

For expression in of YFP-LIN-45 in VPCs, coding sequences for *yfp* and *lin-45* (Wormbase sequence Y73B6A.5a) were fused in frame to produce a sequence encoding N-terminally tagged YFP-LIN-45 protein, and cloned in the vector pCC249 (DE LA COVA AND GREENWALD 2012) or pCC395 (TOWNLEY *et al*. 2023). pCC249 contains miniMos transposon elements and a Neomycin resistance gene derived from pCFJ910 (FROKJAER-JENSEN *et al*. 2014), promoter and regulatory elements of the *lin-31* gene used for VPC-specific expression (MYERS AND GREENWALD 2005), and a 3′ UTR derived from the *unc-54* gene. pCC395 is identical to pCC249 in regulatory sequences, but includes the pharynx-expressed marker *myo-2p*::*tag-RFP* for use in visual selection of transgenic animals.

Single-copy insertion transgenes were generated by germline injection of N2 strain hermaphrodites with 10 ng/μL of the transgene plasmid with a co-injection mixture of pCFJ601, pCFJ90, pGH8, and pMA122. Strains carrying integrated reporters were isolated using the methods of (FROKJAER-JENSEN *et al*. 2014). Briefly, transformant progeny of injected hermaphrodites were selected using G418 (2.5 µg/mL). Surviving progeny without visible co-injection markers were cloned and screened for transgene integration based on Neomycin resistance, RFP expression in the pharynx, or YFP expression in the VPCs.

### Assessment of the Multivulva phenotype

To synchronize adult animals scored for the Multivulva phenotype, hermaphrodites at the L4 larval stage were picked from uncrowded cultures and grown for 24 hours at 20°C. Adults were scored for the presence of pseudovulvae using a dissecting microscope. Animals that had one or more ectopic pseudovulvae were reported as displaying the Multivulva phenotype.

### Statistical analyses

All statistical analyses were performed using GraphPad Prism software. To assess whether multiple, independent transgenic strains carrying the same construct displayed a consistent phenotype, we used counts of Muv and non-Muv animals and performed a Chi-squared test of homogeneity. To make pairwise comparisons between strains carrying different constructs, we used counts of Muv and non-Muv animals and performed a Fisher’s exact test.

### Protein sequence alignments

Protein sequences were aligned using Clustal Omega in Uniprot with default settings (UNIPROT 2023). Alignment figures were produced using Espript (ROBERT AND GOUET 2014). Invertebrate sequences in alignments in Figures S1, S2, and S3 were selected from the set of orthologs identified by EnsemblMetazoa for the mosquito *Anopheles gambiae* Raf (Ensembl Transcript ID AGAP004699-RA), and from orthologs identified by ParaSite for *C. elegans* LIN-45 (Wormbase gene Y73B6A.5), plus the orthologs identified by Uniprot for human BRAF (Uniprot P15056). We used mosquito Raf because it recovered a broader variety of invertebrate sequences than queries with LIN-45.

### YFP-LIN-45 fluorescence quantitation

To synchronize larvae for imaging of VPCs during development, adults were allowed to lay eggs for 2 hours at 25°C and larvae developed for 26-28 hours at 25°C, a time in the L2 larval stage at which LIN-45 protein is not yet subject to protein degradation in P6.p (TOWNLEY *et al*. 2023) Larvae were mounted on an agarose pad in M9 buffer containing 10 mM levamisole. Images used for YFP intensity quantitation were acquired using a Nikon Ti inverted microscope equipped with a spinning disk confocal system (Crest Optics) and sCMOS camera (Teledyne Photometrics). For all images, Z-stacks were acquired using a 40X objective, a 518 nm laser power of 25%, and an exposure time of 700 ms. Image segmentation was performed in NIS-Elements to create regions of interest for the cytoplasm of VPCs. Because LIN-45 protein degradation does not appear to occur in P5.p and P7.p (TOWNLEY *et al*. 2023), we report the mean YFP intensity for the per animal for only these two cells.

### Use of AlphaFold 3 to predict quaternary structure of LIN-45 complexes

Protein structures were predicted using AlphaFold 3 (https://alphafoldserver.com), which permits prediction of multiprotein complexes that contain post-translational modifications, ions, or RNA (ABRAMSON *et al*. 2024). AlphaFold predictions were interpreted using guidance from the EMBL-EBI training manual (https://www.ebi.ac.uk/training/online/courses/alphafold/). ChimeraX was used to open the AlphaFold models (.cif files), visualize the confidence scores (.json files), and manipulate their display to generate figures (MENG *et al*. 2023). To visualize the location of ATP in the kinase domain, the structure of BRAF with the ATP analog AMP-PCP from PDB 6U2G was superimposed with AlphaFold predictions (LIAU *et al*. 2020b). The output from an AlphaFold 3 query produces an ensemble of five models. Confidence can be assessed through the degree of agreement between the ensemble models, and through a residue-by-residue measure of the confidence expressed as pLDDT, a score ranging from 0 to 100.

Our initial AlphaFold queries consisted of two copies of full-length LIN-45, each with the phosphorylation at S756 needed for 14-3-3 binding, and two copies of the *C. elegans* 14-3-3 zeta ortholog PAR-5. For these queries, the regions of LIN-45 and PAR-5 where the ensemble of models were in good agreement were also highly similar to the cryo-EM structures of human BRAF1 and 14-3-3 zeta (PDB 6Q0K) (PARK *et al*. 2019). However, these AlphaFold predictions of LIN-45 had regions with low confidence that did not conform, including novel interactions between the CRD domain and kinase domain, and previously unobserved regions of alpha-helical secondary structure. We reasoned that low confidence interactions could produce clashes that block quaternary structures involving the DTS and kinase domain. Therefore, we removed the N-terminal sequences from LIN-45 in subsequent modeling, using solely the kinase domain and DTS, residues 470-813. We feel this was appropriate as all cryo-EM structures of BRAF reported lack the N-terminal domains (KONDO *et al*. 2019; PARK *et al*. 2019; LIAU *et al*. 2020b; MARTINEZ FIESCO *et al*. 2022).

We modeled LIN-45 complexes with or without phosphorylation at T626 and T629 of the activation loop, and with or without phosphorylation at T791 of the DTS. All models of LIN-45 included phosphorylation at the 14-3-3 binding motif residue S756. For models of complexes with BRAF isoform 1 (NCBI Accession: NP_004324.2) and BRAF isoform 2 (NCBI Accession: NP_001341538.1), we used queries with residues 448-766 for isoform 1 and 448-767 for isoform 2, phosphorylation at the C-terminal 14-3-3 binding site, phosphorylation at S750 and T753 of the DTS, and 14-3-3 zeta (NCBI Accession: NP_663723.1). For *Drosophila melanogaster* Raf complexes, we used queries with Raf (Uniprot KRAF P11346) residues 419-739, phosphorylation at the C-terminal 14-3-3 binding site, phosphorylation at S666 and T669 of the DTS, and 14-3-3 zeta (Uniprot P29310).

## Data availability statement

Strains and plasmids are available upon request. The authors affirm that all data necessary for confirming the conclusions of the article are present within the article, figures, and tables.

## Acknowledgements

Research reported in this publication was supported by the National Cancer Institute under award R03CA248684. The content is solely the responsibility of the authors and does not necessarily represent the official views of the National Institutes of Health. The authors would like to acknowledge the University of Wisconsin-Milwaukee Graduate School for support. We thank Wormbase for *C. elegans* genome and curation data. Some *C. elegans* strains were provided by the Caenorhabditis elegans Genome Center, which is funded by NIH Office of Research Molecular Infrastructure Program (P40 OD010440). We thank AlphaFold Server for protein structure prediction. Molecular graphics and analyses were performed with University of California, San Francisco (UCSF) ChimeraX, developed by the Resource for Biocomputing, Visualization, and Informatics at UCSF, with support from NIH (R01 GM129325 and P41 GM103311).

## Supplementary Materials

Figures S1-S7

Table S1

**Figure S1.**
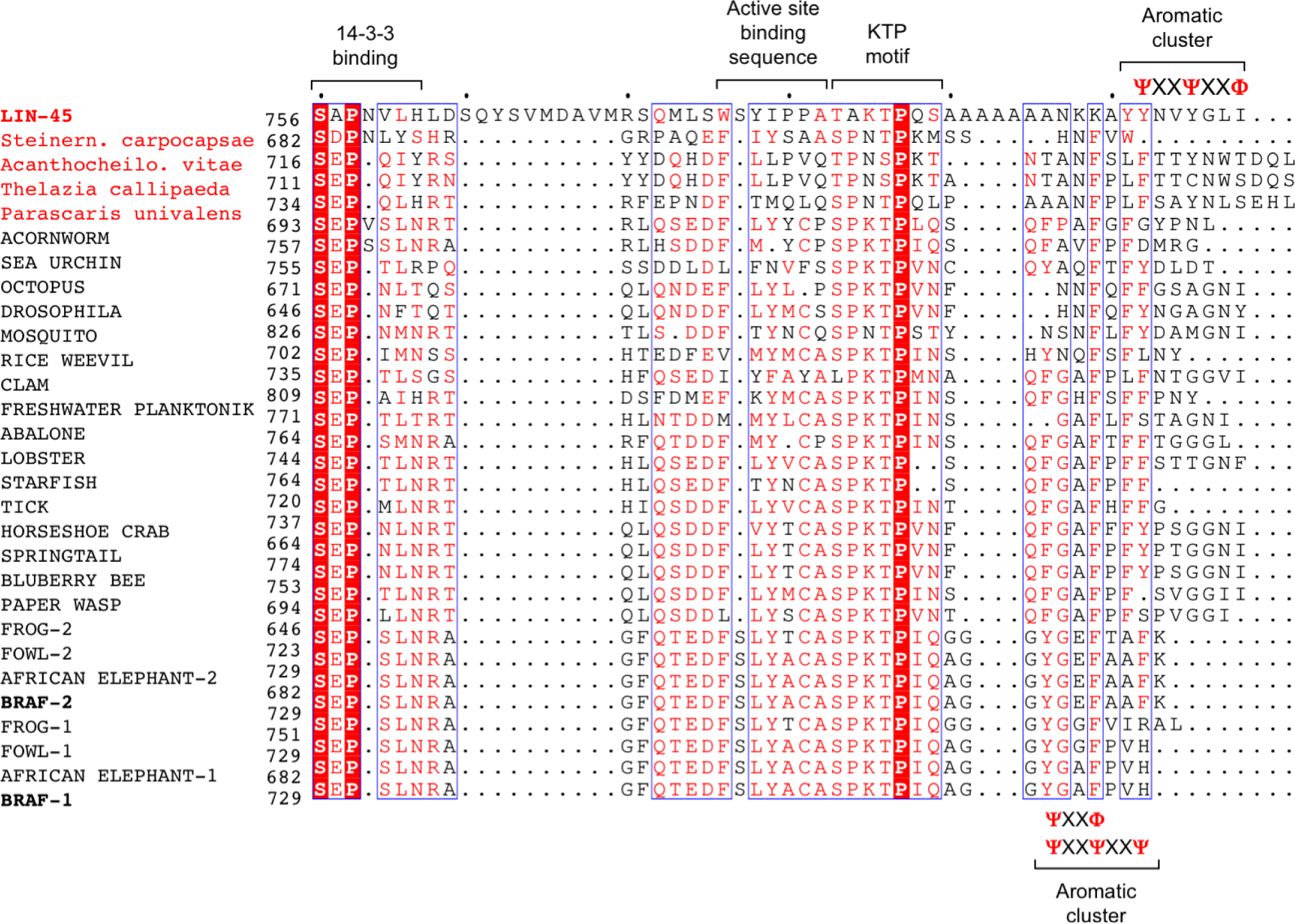
LIN-45 and human BRAF distal tail segment regions share sequence similarity across animals. Alignment of DTS sequences from diverse metazoans. Organisms in red are nematodes. Within the aromatic cluster, possible motifs are indicated. Ψ indicates amino acids Y or F. Φ indicates amino acids F, I, L, M, or V. X indicates any amino acid.

**Figure S2.**
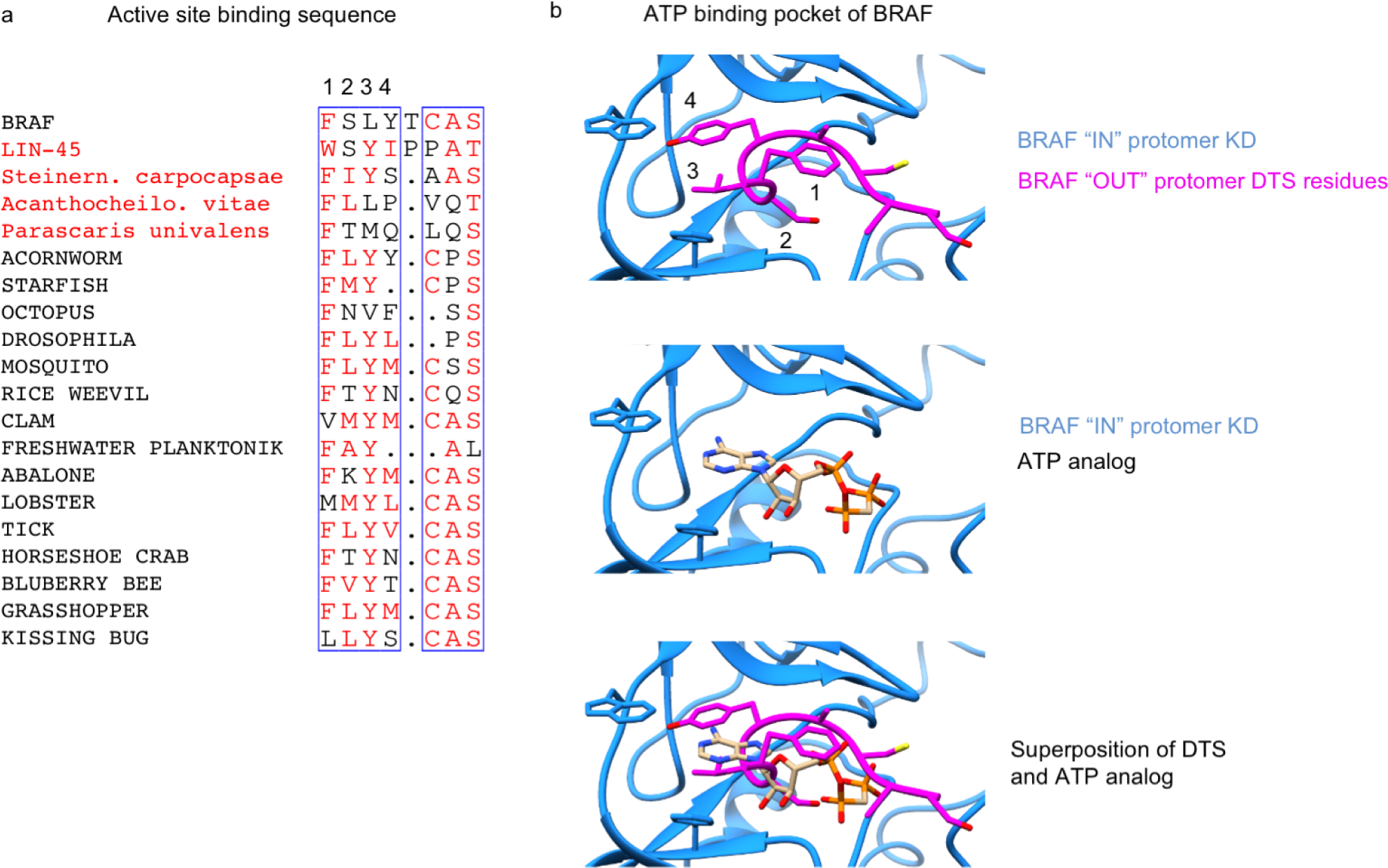
Residues within the BRAF active site binding sequence share common amino acid characteristics with aligned sequences from diverse metazoan. a) A subset of sequences from Fig. S1 within the DTS of Raf proteins from diverse metazoa curated to eliminate redundant sequences. Numbers 1, 2, 3, and 4 labeled above the column indicate position. b) (Top) human BRAF dimer model (PDB 6UAN) with a kinase domain of BRAF “IN” protomer (blue) and DTS of BRAF “OUT” protomer (magenta). Numbers 1, 2, 3, and 4 correspond to positions in the alignment in part a. (Middle) Superposition of kinase domain of BRAF “IN” protomer (blue) with the ATP analog AMP-PCP from model (6U2G). (Bottom) Superposition of kinase domain of BRAF “IN” protomer (blue) and DTS of BRAF “OUT” protomer (magenta) with the ATP analog AMP-PCP.

**Figure S3.**
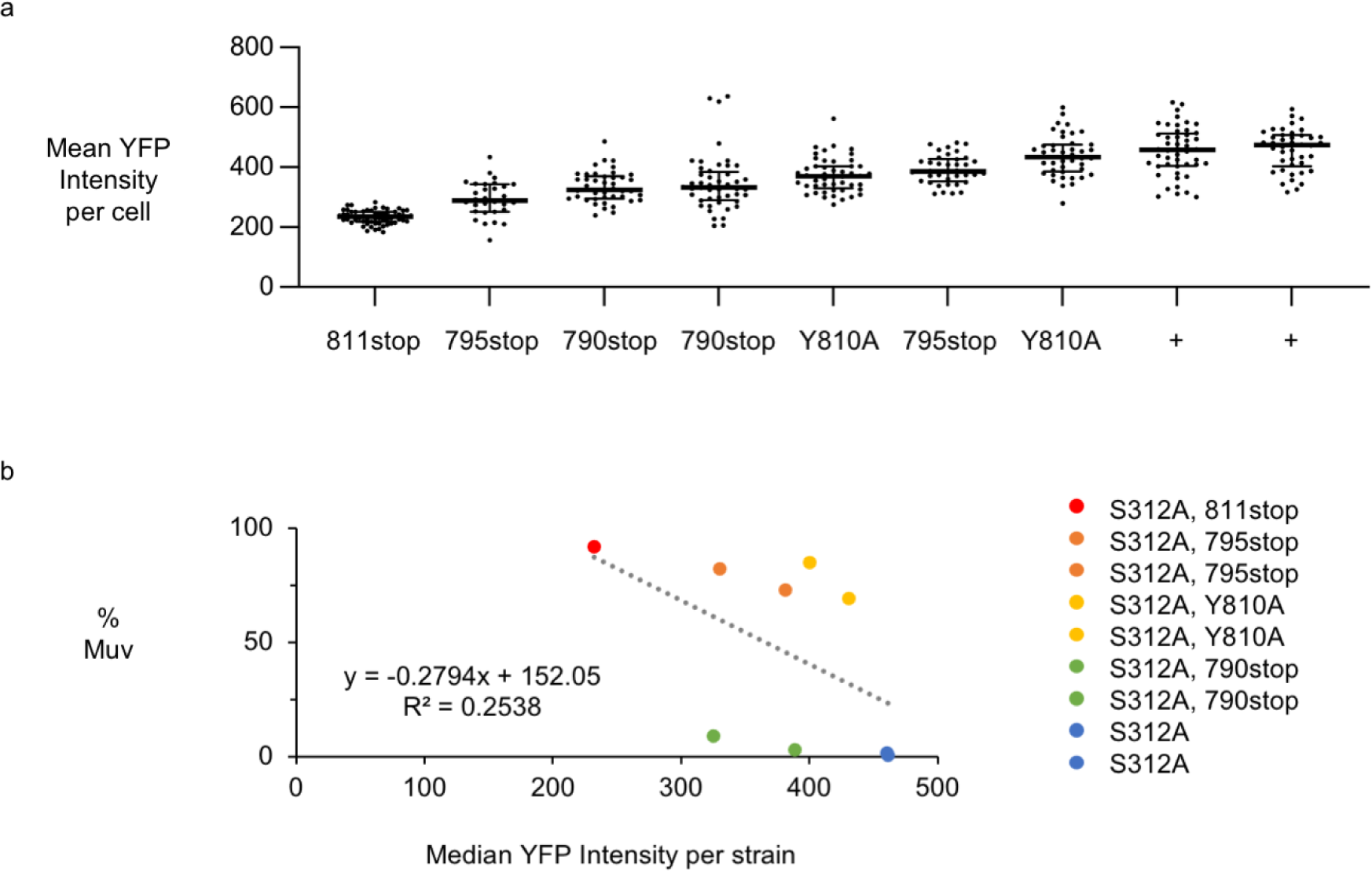
YFP fluorescence intensity in VPCs of mutant *yfp-lin-45(S312A)* transgenic strains. a) Mean YFP fluorescence intensity per cell in the P5.p and P7.p cells of L2 stage larvae. All transgenes encoded YFP-LIN-45(S312A). Additional mutations in each transgene assayed are indicated on the x-axis. b) Penetrance of the Muv phenotype versus median YFP intensity in P5.p and P7.p per strain.

**Figure S4.**
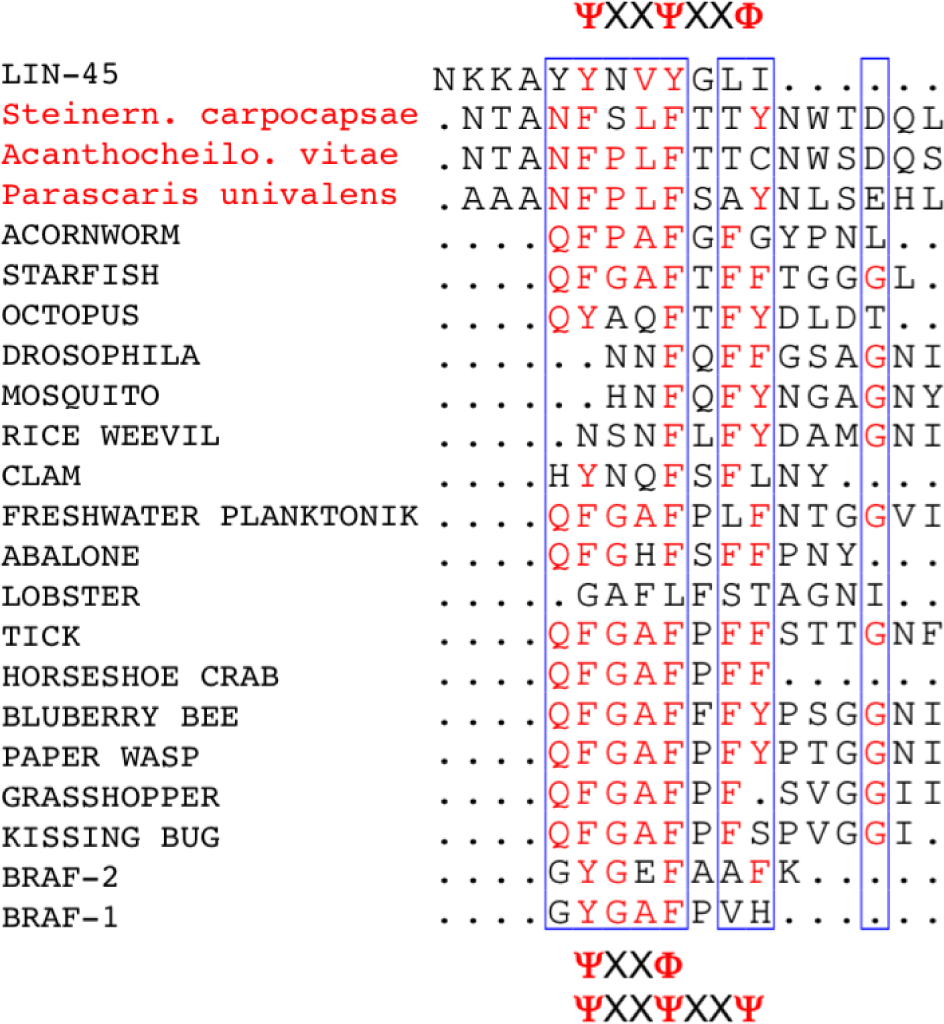
The C-terminus of LIN-45 shares amino acid features with BRAF from diverse metazoans. Manual alignment of LIN-45 aromatic cluster to the curated non redundant sequence set in S2. Possible aromatic cluster motifs are indicated. Ψ indicates amino acids Y or F. Φ indicates amino acids F, I, L, M, or V. X indicates any amino acid.

**Figure S5.**
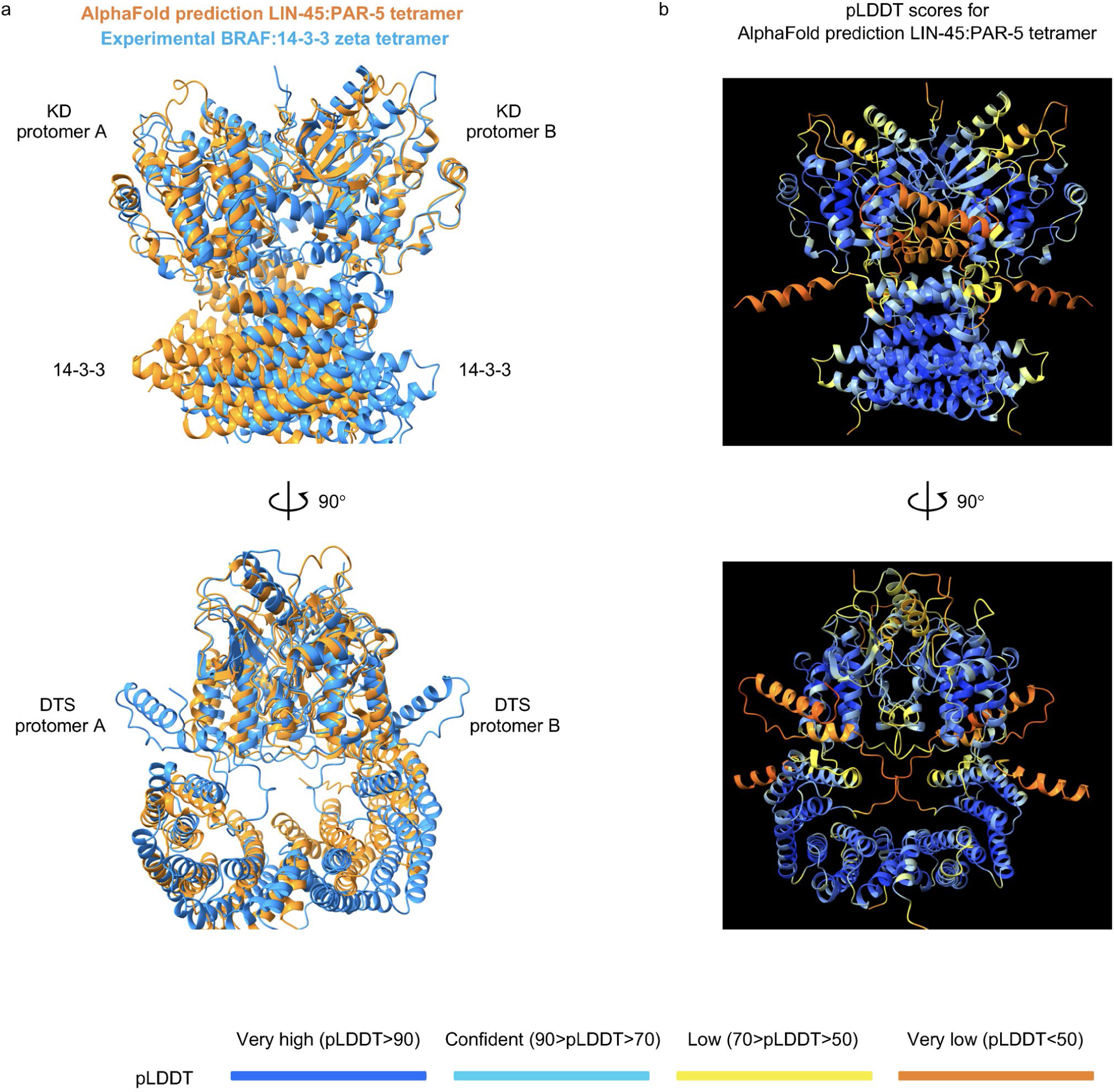
AlphaFold predicts LIN-45:PAR-5 quaternary structure in good agreement with experiment. a) Superposition of the AlphaFold predicted structure of tetrameric LIN-45:PAR-5 (blue) and model of human BRAF:14-3-3 (orange) from PDB 6Q0K. PAR-5 C-termini residues 230-248 have been trimmed for clarity. b) LIN-45:PAR-5 Alphafold prediction confidence pLDDT scores shown as colors, very high (dark blue), confident (blue), low (yellow), very low (orange). PAR-5 model shown without trimming the C-termini.

**Figure S6.**
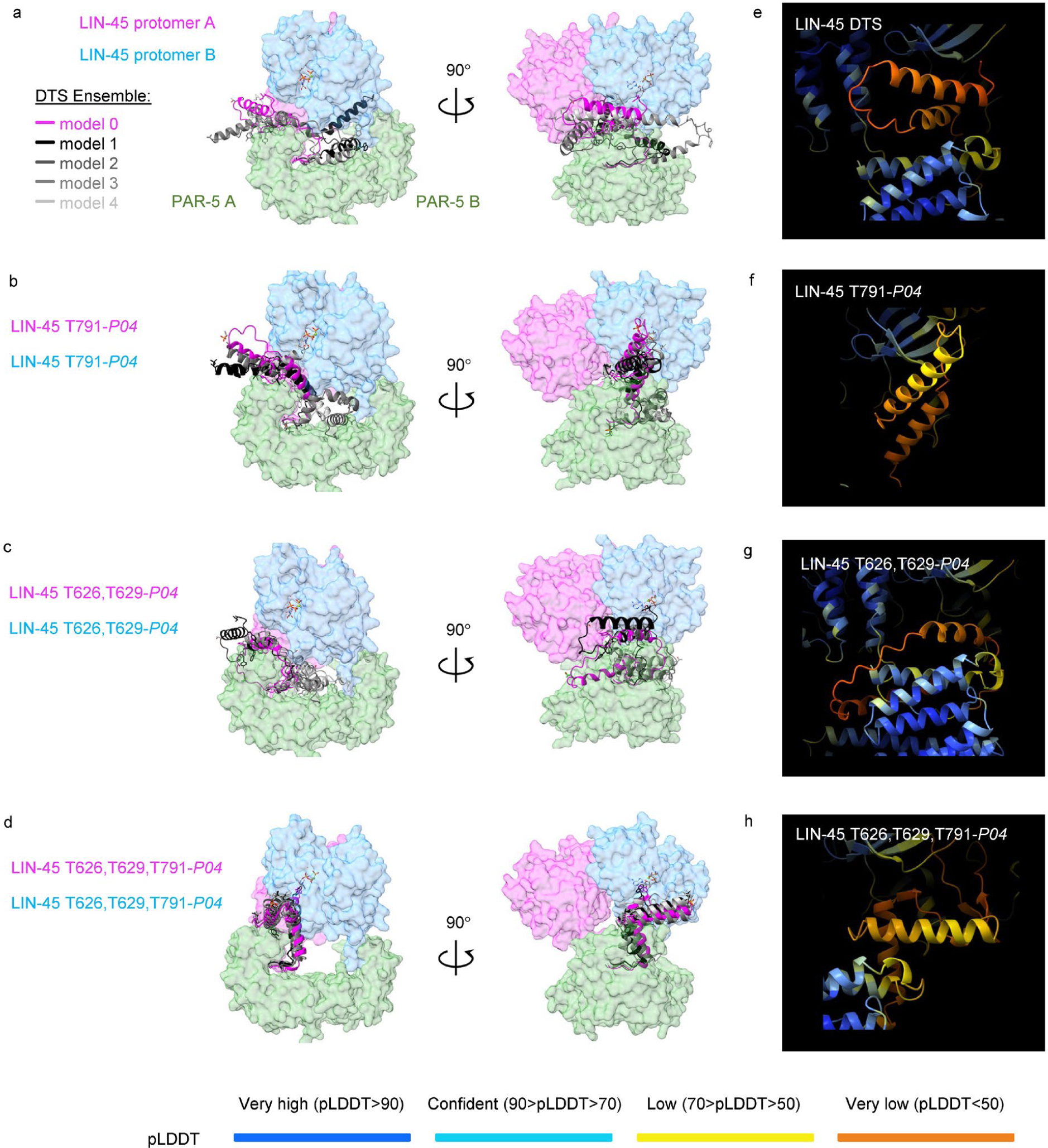
AlphaFold prediction ensemble agreement and confidence scores of the LIN-45 DTS are influenced by activation loop and T791 phosphorylation. a-d) All panels show two orthogonal views of AlphaFold predicted tetramer containing LIN-45 protomers A (magenta) and B (blue) and PAR-5 protomers A and B (green). All predictions included phosphorylation at the S756 site needed for 14-3-3 binding. The two kinase domains and PAR-5 proteins are depicted as surfaces. For the DTS, all five ensemble models are superimposed and depicted as ribbons (magenta, black, dark gray, medium gray, light gray). PAR-5 C-termini residues 230-248 have been trimmed for clarity. a) LIN-45. b) LIN-45 with pT791. c) LIN-45 with pT626 and pT629. d) LIN-45 with pT626, pT629, T791. e-h) All panels show Alphafold prediction confidence pLDDT scores for the LIN-45 DTS depicted as colors. e) LIN-45. f) LIN-45 with pT791. g) LIN-45 with pT626 and pT629. h) LIN-45 with pT626, pT629, T791.

**Figure S7.**
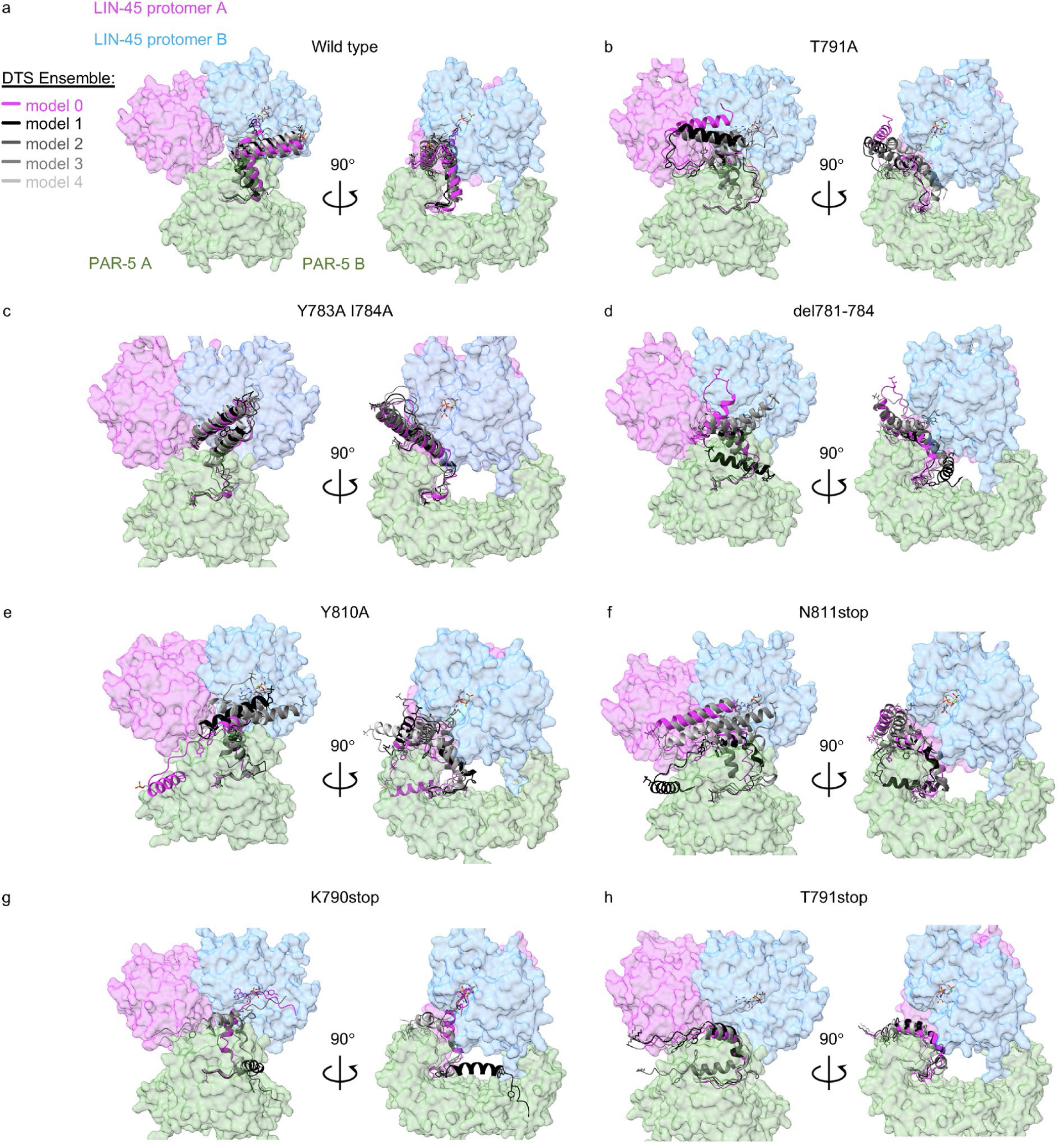
AlphaFold prediction of LIN-5 structural changes resulting from DTS mutations. All panels show two orthogonal views of AlphaFold predicted tetramer containing LIN-45 protomers A (magenta) and B (blue) and PAR-5 protomers A and B (green). All predictions included phosphorylation at the S756 site needed for 14-3-3 binding, the activation loop (pT626, pT629), and DTS if present (pT791). The two kinase domains and PAR-5 proteins are depicted as surfaces. For the DTS, all five ensemble models are superimposed and depicted as ribbons (magenta, black, dark gray, medium gray, light gray). PAR-5 C-termini residues 230-248 have been trimmed for clarity. a) Wild-type LIN-45. b-h) Mutant LIN-45 forms shown are: b) T791A, c) Y783A, I784A, d) del781-784, e) Y810A, e) N811stop, f) K790stop, and g) T791stop.

**Table S1.** *C. elegans* strains used in this work. For each strain, the corresponding data figure, strain name, genotype, number of animals scored (n), count of non-Muv, count of Muv, and percentage of Muv are listed.

| Fig | Strain | Genotype | N | non-Muv | Muv | % Muv |
| --- | --- | --- | --- | --- | --- | --- |
| 2b | MKE212 | *covTi62 [lin-31p::yfp-lin-45::unc-54 3'UTR, myo-2p::tagRFP::unc-54 3'UTR]* | 80 | 80 | 0 | 0 |
| 2b | MKE214 | *covTi64 [lin-31p::yfp-lin-45::unc-54 3'UTR, myo-2p::tagRFP::unc-54 3'UTR]* | 290 | 290 | 0 | 0 |
| 2b | MKE298 | *covTi97 [lin-31p::yfp-lin-45(S312A)::unc-54 3'UTR, myo-2p::tagRFP::unc-54 3'UTR]* | 399 | 395 | 4 | 1 |
| 2b | MKE299 | *covTi98 [lin-31p::yfp-lin-45(S312A)::unc-54 3'UTR, myo-2p::tagRFP::unc-54 3'UTR]* | 515 | 512 | 3 | 1 |
| 2b | MKE300 | *covTi99 [lin-31p::yfp-lin-45(S312A)::unc-54 3'UTR, myo-2p::tagRFP::unc-54 3'UTR]* | 450 | 446 | 4 | 1 |
| 2b | MKE302 | *covTi101 [lin-31p::yfp-lin-45(S312A)::unc-54 3'UTR, myo-2p::tagRFP::unc-54 3'UTR]* | 621 | 610 | 11 | 2 |
| 2c | MKE426 | *covTi127 [lin-31p::yfp-lin-45(773stop)::unc-54 3'UTR, myo-2p::tagRFP::unc-54 3'UTR]* | 94 | 94 | 0 | 0 |
| 2c | MKE352 | *covTi117 [lin-31p::yfp-lin-45(763stop)::unc-54 3'UTR, myo-2p::tagRFP::unc-54 3'UTR]* | 62 | 62 | 0 | 0 |
| 2c | MKE489 | *covTi172 [lin-31p::yfp-lin-45(312A,773stop)::unc-54 3'UTR, myo-2p::tagRFP::unc-54 3'UTR]* | 59 | 5 | 54 | 92 |
| 2c | MKE490 | *covTi173 [lin-31p::yfp-lin-45(312A,773stop)::unc-54 3'UTR, myo-2p::tagRFP::unc-54 3'UTR]* | 41 | 1 | 40 | 98 |
| 2c | MKE356 | *covTi119 [lin-31p::yfp-lin-45(312A,763stop)::unc-54 3'UTR, myo-2p::tagRFP::unc-54 3'UTR]* | 104 | 0 | 104 | 100 |
| 2c | MKE453 | *covTi119 [lin-31p::yfp-lin-45(312A,763stop)::unc-54 3'UTR, myo-2p::tagRFP::unc-54 3'UTR]; lin-45(dx19)/nT1* | 21 | 0 | 21 | 100 |
| 2d | MKE469 | *covTi163 [lin-31p::yfp-lin-45(175E,177E)::unc-54 3'UTR, myo-2p::tagRFP::unc-54 3'UTR]* | 90 | 0 | 0 | 0 |
| 2d | MKE606 | *covTi260 [lin-31p::yfp-lin-45(175E,177E,773stop)::unc-54 3'UTR]* | 60 | 0 | 0 | 0 |
| 2d | MKE634 | *covTi249 [lin-31p::yfp-lin-45(T177P)::unc-54 3'UTR]* | 100 | 0 | 0 | 0 |
| 2d | MKE620 | *covTi268 [lin-31p::yfp-lin-45(T177P,773stop)::unc-54 3'UTR]* | 110 | 0 | 0 | 0 |
| 2e | MKE498 | *covTi163 [lin-31p::yfp-lin-45(175E,177E)::unc-54 3'UTR, myo-2p::tagRFP::unc-54 3'UTR]; lin-45(dx19)/nT1* | 65 | 46 | 19 | 29 |
| 2e | MKE1001 | *covTi260 [lin-31p::yfp-lin-45(175E,177E,773stop)::unc-54 3'UTR]; lin-45(dx19)/nT1* | 48 | 42 | 6 | 13 |
| 2e | MKE997 | *covTi249 [lin-31p::yfp-lin-45(T177P)::unc-54 3'UTR]; lin-45(dx19)/nT1* | 8 | 8 | 0 | 0 |
| 2e | MKE1000 | *covTi268 [lin-31p::yfp-lin-45(T177P,773stop)::unc-54 3'UTR]; lin-45(dx19)/nT1* | 21 | 21 | 0 | 0 |
| 3c | MKE494 | *covTi174 [lin-31p::yfp-lin-45(S312A,781stop)::unc-54 3'UTR]* | 37 | 0 | 37 | 100 |
| 3c | MKE529 | *covTi200 [lin-31p::yfp-lin-45(S312A,790stop)::unc-54 3'UTR]* | 92 | 89 | 3 | 3 |
| 3c | MKE530 | *covTi201 [lin-31p::yfp-lin-45(S312A,790stop)::unc-54 3'UTR]* | 106 | 96 | 10 | 9 |
| 3c | MKE531 | *covTi202 [lin-31p::yfp-lin-45(S312A,790stop)::unc-54 3'UTR]* | 110 | 103 | 7 | 6 |
| 3c | MKE532 | covTi203 [lin-31p::yfp-lin-45(S312A,790stop)::unc-54 3'UTR] | 103 | 100 | 3 | 3 |
| 3c | MKE574 | covTi228 [lin-31p::yfp-lin-45(S312A,791stop)::unc-54 3'UTR] | 179 | 40 | 139 | 78 |
| 3c | MKE575 | covTi229 [lin-31p::yfp-lin-45(S312A,791stop)::unc-54 3'UTR] | 54 | 18 | 36 | 67 |
| 3c | MKE576 | covTi230 [lin-31p::yfp-lin-45(S312A,791stop)::unc-54 3'UTR] | 126 | 32 | 94 | 75 |
| 3c | MKE534 | covTi204 [lin-31p::yfp-lin-45(S312A,795stop)::unc-54 3'UTR] | 65 | 42 | 23 | 35 |
| 3c | MKE536 | covTi206 [lin-31p::yfp-lin-45(S312A,795stop)::unc-54 3'UTR] | 121 | 31 | 90 | 74 |
| 3c | MKE537 | covTi207 [lin-31p::yfp-lin-45(S312A,795stop)::unc-54 3'UTR] | 106 | 11 | 95 | 90 |
| 3c | MKE538 | covTi208 [lin-31p::yfp-lin-45(S312A,795stop)::unc-54 3'UTR] | 97 | 27 | 70 | 72 |
| 3c | MKE539 | covTi209 [lin-31p::yfp-lin-45(S312A,795stop)::unc-54 3'UTR] | 57 | 10 | 47 | 83 |
| 3d | MKE753 | covTi326 [lin-31p::yfp-lin-45(S312A,783A,784A)::unc-54 3'UTR] | 774 | 751 | 23 | 3 |
| 3d | MKE754 | covTi327 [lin-31p::yfp-lin-45(S312A,783A,784A)::unc-54 3'UTR] | 484 | 446 | 38 | 8 |
| 3d | MKE761 | covTi329 [lin-31p::yfp-lin-45(S312A,783A,784A)::unc-54 3'UTR] | 592 | 557 | 35 | 6 |
| 3d | MKE869 | covTi412 [lin-31p::yfp-lin-45(S312A,del781-784)::unc-54 3'UTR] | 111 | 110 | 1 | 1 |
| 3d | MKE870 | covTi413 [lin-31p::yfp-lin-45(S312A,del781-784)::unc-54 3'UTR] | 228 | 217 | 11 | 5 |
| 3d | MKE871 | covTi414 [lin-31p::yfp-lin-45(S312A,del781-784)::unc-54 3'UTR] | 113 | 110 | 3 | 3 |
| 3d | MKE559 | covTi211 [lin-31p::yfp-lin-45(S312A,T791A)::unc-54 3'UTR] | 91 | 89 | 2 | 2 |
| 3d | MKE561 | covTi212 [lin-31p::yfp-lin-45(S312A,T791A)::unc-54 3'UTR] | 105 | 100 | 5 | 5 |
| 3e | MKE302 | covTi101 [lin-31p::yfp-lin-45(S312A)::unc-54 3'UTR, myo-2p::tagRFP::unc-54 3'UTR]; lin-45(dx19)/nT1 | 106 | 94 | 12 | 11 |
| 3e | MKE964 | covTi327 [lin-31p::yfp-lin-45(S312A,783A,784A)::unc-54 3'UTR]; lin-45(dx19)/nT1 | 46 | 23 | 23 | 50 |
| 3e | MKE965 | covTi329 [lin-31p::yfp-lin-45(S312A,783A,784A)::unc-54 3'UTR]; lin-45(dx19)/nT1 | 43 | 25 | 18 | 42 |
| 3e | MKE899 | covTi413 [lin-31p::yfp-lin-45(S312A,del781-784)::unc-54 3'UTR]; lin-45(dx19)/nT1 | 84 | 53 | 31 | 37 |
| 3e | MKE1002 | covTi211 [lin-31p::yfp-lin-45(S312A,T791A)::unc-54 3'UTR]; lin-45(dx19)/nT1 | 243 | 170 | 73 | 30 |
| 3e | MKE963 | covTi201 [lin-31p::yfp-lin-45(S312A,790stop)::unc-54 3'UTR]; lin-45(dx19)/nT1 | 37 | 22 | 15 | 41 |
| 4a | MKE737 | covTi314 [lin-31p::yfp-lin-45(S312A,808stop)::unc-54 3'UTR] | 101 | 46 | 55 | 55 |
| 4a | MKE738 | covTi315 [lin-31p::yfp-lin-45(S312A,808stop)::unc-54 3'UTR] | 97 | 13 | 84 | 87 |
| 4a | MKE739 | covTi316 [lin-31p::yfp-lin-45(S312A,808stop)::unc-54 3'UTR] | 93 | 13 | 80 | 86 |
| 4a | MKE815 | covTi364 [lin-31p::yfp-lin-45(S312A,811stop)::unc-54 3'UTR] | 115 | 17 | 98 | 85 |
| 4a | MKE816 | covTi365 [lin-31p::yfp-lin-45(S312A,811stop)::unc-54 3'UTR] | 99 | 53 | 46 | 47 |
| 4a | MKE817 | covTi366 [lin-31p::yfp-lin-45(S312A,811stop)::unc-54 3'UTR] | 103 | 10 | 93 | 90 |
| 4a | MKE818 | covTi367 [lin-31p::yfp-lin-45(S312A,811stop)::unc-54 3'UTR] | 114 | 9 | 105 | 92 |
| 4b | MKE834 | covTi382 [lin-31p::yfp-lin-45(S312A,Y810A)::unc-54 3'UTR] | 105 | 16 | 89 | 85 |
| 4b | MKE837 | covTi385 [lin-31p::yfp-lin-45(S312A,Y810A)::unc-54 3'UTR] | 125 | 39 | 86 | 69 |
| 4b | MKE838 | covTi386 [lin-31p::yfp-lin-45(S312A,Y810A)::unc-54 3'UTR] | 117 | 28 | 89 | 76 |
| 4b | MKE1005 | covTi386 [lin-31p::yfp-lin-45(S312A,Y810A)::unc-54 3'UTR]; lin-45(dx19)/nT1 | 23 | 0 | 23 | 100 |
| 4c | MKE214 | covTi64 [lin-31p::yfp-lin-45::unc-54 3'UTR, myo-2p::tagRFP::unc-54 3'UTR] | 290 | 290 | 0 | 0 |
| 4c | MKE917 | covTi64 [lin-31p::yfp-lin-45::unc-54 3'UTR, myo-2p::tagRFP::unc-54 3'UTR]; dpy-23(e840) | 155 | 143 | 12 | 8 |
| 4c | MKE895 | covTi64 [lin-31p::yfp-lin-45::unc-54 3'UTR, myo-2p::tagRFP::unc-54 3'UTR]; gap-1(ga133) | 143 | 142 | 1 | 1 |
| 4c | MKE298 | covTi97 [lin-31p::yfp-lin-45(S312A)::unc-54 3'UTR, myo-2p::tagRFP::unc-54 3'UTR] | 399 | 395 | 4 | 1 |
| 4c | MKE918 | covTi97 [lin-31p::yfp-lin-45(S312A)::unc-54 3'UTR, myo-2p::tagRFP::unc-54 3'UTR]; dpy-23(e840) | 122 | 80 | 42 | 34 |
| 4c | MKE889 | covTi197 [lin-31p::yfp-lin-45(S312A)::unc-54 3'UTR, myo-2p::tagRFP::unc-54 3'UTR]; gap-1(ga133) | 108 | 4 | 104 | 96 |
| 4c | MKE302 | covTi101 [lin-31p::yfp-lin-45(S312A)::unc-54 3'UTR, myo-2p::tagRFP::unc-54 3'UTR] | 621 | 610 | 11 | 2 |
| 4c | MKE919 | covTi101 [lin-31p::yfp-lin-45(S312A)::unc-54 3'UTR, myo-2p::tagRFP::unc-54 3'UTR]; dpy-23(e840) | 222 | 141 | 81 | 36 |
| 4c | MKE886 | covTi101 [lin-31p::yfp-lin-45(S312A)::unc-54 3'UTR, myo-2p::tagRFP::unc-54 3'UTR]; gap-1(ga133) | 75 | 1 | 74 | 99 |
| 4c | MKE530 | covTi201 [lin-31p::yfp-lin-45(S312A,790stop)::unc-54 3'UTR] | 106 | 96 | 10 | 9 |
| 4c | MKE924 | covTi201 [lin-31p::yfp-lin-45(S312A,790stop)::unc-54 3'UTR]; dpy-23(e840) | 180 | 113 | 67 | 37 |
| 4c | MKE995 | covTi201 [lin-31p::yfp-lin-45(S312A,790stop)::unc-54 3'UTR]; gap-1(ga133) | 104 | 11 | 93 | 89 |
| 4d | MKE937 | covTi438 [lin-31p::yfp-lin-45(Q95A,R119A,S312A,Y810A)::unc-54 3'UTR] | 211 | 210 | 1 | 0 |
| 4d | MKE938 | covTi439 [lin-31p::yfp-lin-45(Q95A,R119A,S312A,Y810A)::unc-54 3'UTR] | 100 | 100 | 0 | 0 |
| 4d | MKE939 | covTi440 [lin-31p::yfp-lin-45(Q95A,R119A,S312A,Y810A)::unc-54 3'UTR] | 161 | 161 | 0 | 0 |
| 4d | MKE956 | covTi449 [lin-31p::yfp-lin-45(Q95A,R119A,V627E)::unc-54 3'UTR] | 94 | 94 | 0 | 100 |
| 4d | MKE957 | covTi451 [lin-31p::yfp-lin-45(Q95A,R119A,V627E)::unc-54 3'UTR] | 54 | 1 | 53 | 98 |
| 4d | MKE959 | covTi452 [lin-31p::yfp-lin-45(Q95A,R119A,V627E)::unc-54 3'UTR] | 72 | 3 | 69 | 96 |
| 4e | RB915 | ksr-1(ok786) | 100 | 100 | 0 | 0 |
| 4e | MKE537 | covTi207 [lin-31p::yfp-lin-45(S312A,795stop)::unc-54 3'UTR] | 90 | 5 | 85 | 94 |
| 4e | MKE675 | covTi207 [lin-31p::yfp-lin-45(S312A,795stop)::unc-54 3'UTR]; ksr-1(ok786) | 86 | 85 | 1 | 1 |

